# Spatiotemporal diversification of forest understorey species reveals the existence of multiple Pleistocene forest refugia in Central Europe

**DOI:** 10.1101/2025.05.13.653645

**Authors:** Camille Voisin, Philipp Kirschner, Eliška Záveská, Božo Frajman, Karl Hülber, Johannes Wessely, Wolfgang Willner, Peter Schönswetter, Pau Carnicero

**Affiliations:** Department of Botany, University of Innsbruck, Innsbruck, Austria; Laboratoire d’Écologie Alpine, Université Grenoble Alpes, Université Savoie Mont Blanc, CNRS, Grenoble, France; Institute of Botany of the Czech Academy of Sciences, Průhonice, Czechia; Department of Botany and Biodiversity Research, University of Vienna, Vienna, Austria; Vienna Institute for Nature Conservation & Analyses (VINCA), Vienna, Austria; Department of Animal Biology, Plant Biology and Ecology, Autonomous University of Barcelona, Bellaterra, Spain

**Keywords:** forest understorey species, demographic modelling, glacial refugia, phylogeography, RAD sequencing

## Abstract

During Pleistocene cold stages, European temperate forests were not only restricted to refugia in the southern European peninsulas. Rather, there is increasing evidence for survival of trees also in isolated patches further north, termed “northern refugia”. While their existence is undisputed, based on what is known from a handful of tree species, there is very limited knowledge about forest understoreys. Here, we fill this gap by examining the evolutionary histories of three Central European forest understorey species (FUS; Aposeris foetida, Cardamine trifolia, Hacquetia epipactis). To do so, we use a set of exploratory and explicit analyses utilizing genomic data and ecological niche models, and interpret these data following an a priori defined framework. We identify the northwestern Balkan Peninsula as the primary diversification center for the three species but found additional northern refugia in the Alps, the Carpathians, and the Apennines. Divergence times indicated pre-Last Glacial Maximum (LGM) diversification within each FUS, suggesting persistence of forest islands in Central Europe during Pleistocene cold stages rather than exclusively post-LGM colonization. We conclude that FUS thrived in scattered northern refugia. This refines our understanding of past forest dynamics and further supports widespread long-term persistence of forest patches in Central Europe during Pleistocene cold stages.

## Introduction

The geographic range of temperate forests, which are currently the dominant zonal vegetation of large parts of Europe (Leuschner & Ellenberg, 2017), has been strongly reshuffled by Quaternary climatic fluctuations (Birks & Tinner, 2016; Magri et al., 2006; Petit et al., 2003). Phases of drastic range reductions to small and disjunct refugia during cold stages alternated with phases of expansions leading to continuous distributions during warm stages such as the Holocene (Comes & Kadereit, 1998). Last Glacial Maximum (LGM, ∼ 21 ka) refugia for temperate deciduous trees were initially proposed only in the southern European peninsulas (Hewitt, 1999; Taberlet et al., 1998). However, subsequent studies suggested additional refugia north of these peninsulas, that are often also named ‘northern refugia’ or ‘cryptic refugia’ (Stewart & Lister, 2001; Postolache *et al*., 2017; conceptual discussion in Rull, 2010; hereafter only the term ‘northern refugia’ will be used).

Several northern refugia for temperate forests were proposed for Central Europe, the northern Apennines, the southwesternmost Alps and the southern Carpathians (Fig. 1). Specifically, for European beech (*Fagus sylvatica*), one of the dominant trees of European temperate forests, macrofossil and palynological evidence alongside genetic data suggested northern refugia in the northwestern Balkan Peninsula, the southwestern fringe of the Western Alps, the Western Carpathians and the Apuseni Mountains (Fig. 1; Magri *et al*., 2006; Brus, 2010). Genomic data further supported a northern beech refugium in the Apennines (Marchesini et al., 2023). The existence of an isolated refugium of broad-leaved trees in the Euganean Hills (Colli Euganei) at the southern margin of the Alps (Gubler et al., 2018; Kaltenrieder et al., 2009) and in a former hydrothermal field in Moravia (Hošek et al., 2024) were further suggested by pollen data as well as retrospective evaluation of current microclimatic variability. For coniferous trees, additional refugia were discovered at the southern and south-eastern edge of the Alps and in the adjacent Po Plains (Gugerli et al., 2023; Monegato et al., 2015; van der Knaap et al., 2005), the forelands of the easternmost Central Alps (e.g. Huber *et al*., 2010) and even the easterly adjacent Pannonian Plains (Fig. 1; Willis & van Andel, 2004; Gugerli *et al*., 2023).

**Fig. 1.**
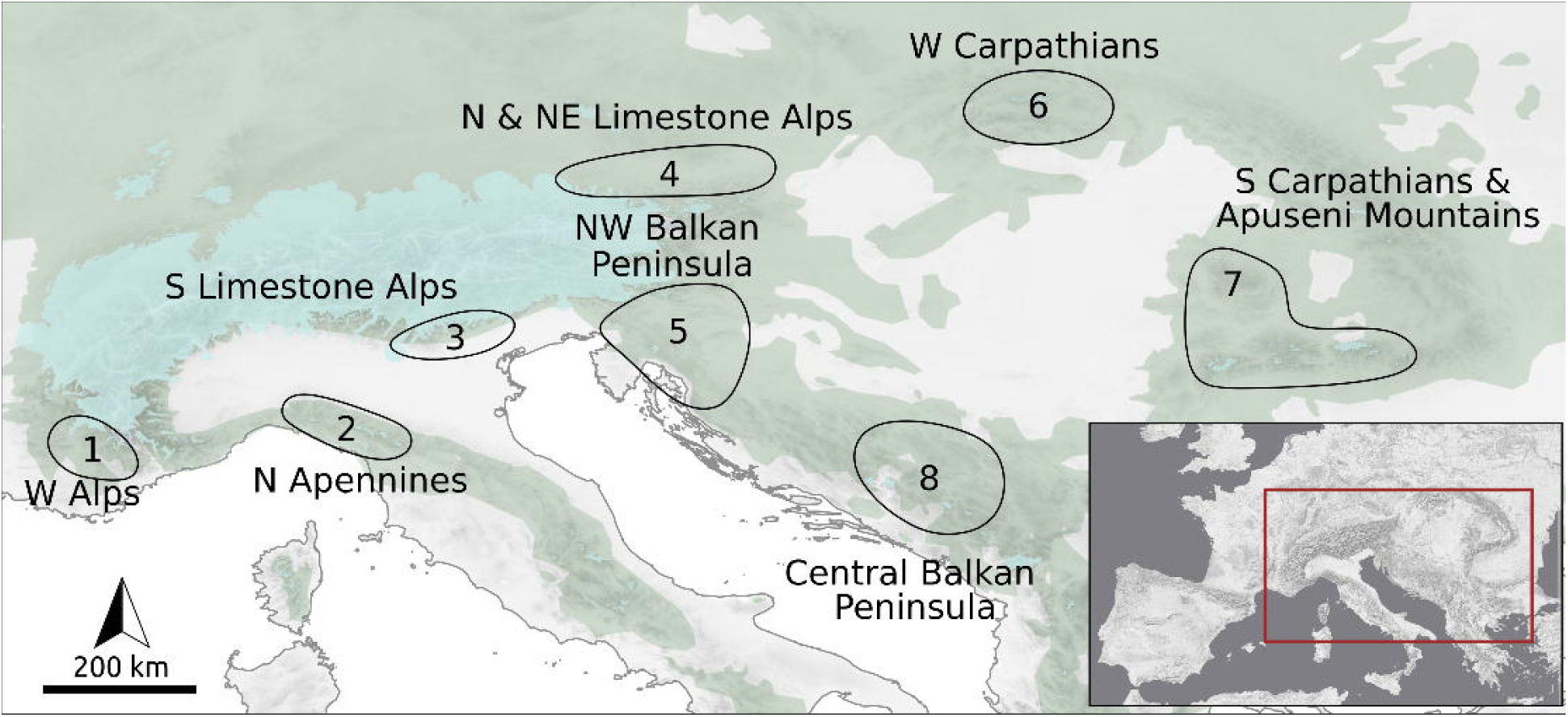
Potential refugia of the studied forest understorey species in Central Europe and neighboring areas during the Last Glacial Maximum (LGM, ∼21 ka). Green areas refer to the current distribution of European beech (taken from(Willner et al., 2023)) and reflect the potential natural distribution of temperate forest. Light blue areas indicate the maximum extent of the LGM glaciation (Ehlers et al., 2011). **1** Western Alps (Magri et al., 2006); **2** northern Apennines (Guido et al., 2020; Marchesini et al., 2023; Willner et al., 2009); **3** Southern Limestone Alps (Brus, 2010; Daneck et al., 2016; Ravazzi et al., 2012; Slovák et al., 2012; Záveská et al., 2021); **4** Northeastern Limestone Alps (Záveská et al., 2021); **5** northwestern Balkan Peninsula (Birks & Willis, 2008; Brus, 2010; Kirschner et al., 2023; Magri et al., 2006; Monegato et al., 2015; Willis & van Andel, 2004; Willner et al., 2009); **6** Western Carpathians (Birks & Willis, 2008; Daneck et al., 2016; Magri et al., 2006; Magyari, 2002; Urbaniak et al., 2018; Willis & van Andel, 2004); **7** Southern Carpathians & Apuseni Mountains (Kirschner et al., 2023; Magri et al., 2006); **8** central Balkan Peninsula (Daneck et al., 2016; Kirschner et al., 2023; Magri et al., 2006; Slovák et al., 2012);

Most dominant European tree species are wind-pollinated, which facilitates extensive pollen-mediated gene flow over long distances, and promotes homogenisation of gene pools (Buiteveld et al., 2007; Comps et al., 2001; Wessinger, 2021). While this ultimately confers blurred genetic signals, especially on smaller spatial scales, the application of a large number of genomic markers has recently refined our knowledge of genetic structure in some wind-pollinated tree species on a continental scale (Milesi et al., 2024). In contrast to trees, most temperate forest understorey species (hereafter FUS) are insect-pollinated herbs with limited pollen and seed dispersal capacity, which implies reduced rates of gene flow and more conserved patterns of genetic variation as compared to trees, especially on smaller spatial scales. Phylogeographic studies on FUS thus refined our understanding of the Pleistocene history of temperate forests in Europe in unparalleled detail (Kirschner et al., 2023; Slovák et al., 2012; Urbaniak et al., 2018; Záveská et al., 2021). Specifically, alongside corroborating refugia inferred for trees, they pointed at the existence of additional forest refugia along the southern margin of the Southern Limestone Alps (Fig. 1; *Cyclamen purpurascens*, Slovák et al., 2012; *Helleborus niger*, Záveská et al., 2021), and in the Northeastern Limestone Alps (*H. niger;* Záveská et al., 2021).

Beyond fossil records and phylogeographic evidence, ecological niche models (ENMs) have emerged as the third major source of evidence to infer areas with climatic stability throughout the glacial-interglacial climate fluctuations of the Quaternary (Gavin et al., 2014). When projected to past climatic conditions, such models allow the integration of ecological data and provide detailed information on environmentally suitable areas under different climatic conditions (Kirchheimer et al., 2018). Specifically, ENMs supported the presence of European beech in northern refugia, such as the Carpathians (Saltré et al., 2013), and also their absence from northern Europe (Svenning, Normand, & Skov, 2008). In contrast to trees, ENMs were rarely used to reconstruct refugia of FUS. Willner *et al*. 2023 modelled dispersal processes to explain the highly idiosyncratic distribution patterns of a set of species including the ones used in this study. They found that the small (relative to those of trees) and fragmented ranges of many FUS are best explained by a combination of broad ecological niches and rare medium- and long-distance dispersal events. Stochasticity is thus an important determinant of current species ranges, explaining the idiosyncratic distribution patterns of the study species despite strong similarities in refugia, ecological tolerances and dispersal abilities.

Approaches combining genomic data and ENMs have been extensively used to provide large-scale inferences about past range shifts and refugial dynamics of species and entire biomes (Carnicero et al., 2022; Kirschner et al., 2020, 2023; Pan et al., 2020; Rota et al., 2024). Genetic and ecological cues commonly interpreted as evidence of refugial survival of populations or lineages are presented in Fig. 2. In essence, alongside pre-LGM divergence followed by strong isolation, such cues rely on the spatial aggregation of private genetic variation and the geographic distribution of distinct genetic lineages (Widmer & Lexer, 2001; Tribsch & Schönswetter, 2003; Paun *et al*., 2008; Šrámková-Fuxová *et al*., 2017). However, a direct translation of such patterns into hypotheses about the number and location of cold-stage refugia would be overly simplistic, as demographic processes such as bottlenecks could ultimately produce very similar patterns of genetic variation in a much shorter time. As a possible solution to this dilemma, demographic modelling based on genomic data allows discriminating between evolutionary scenarios, such as long-term refugial survival and post-LGM foundation of a population, in a rigorous statistical framework (Gutenkunst et al., 2009), providing at the same time deeper insights into these processes (Kirschner et al., 2023; Rota et al., 2024; Záveská et al., 2021).

**Fig. 2.**
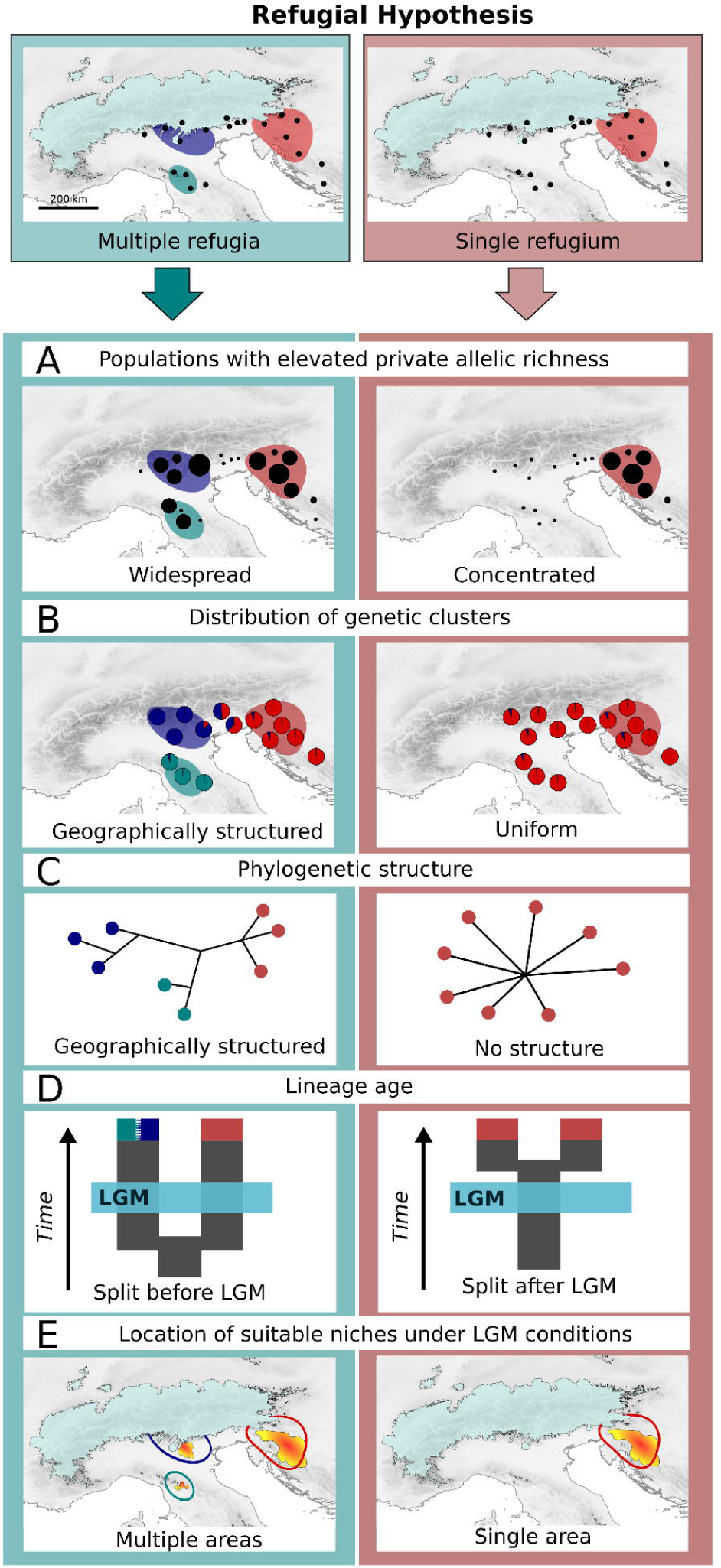
Criteria applied for distinguishing between evolutionary hypotheses involving multiple refugia versus a single refugium for the studied forest understorey species. Black dots refer to the set of sampled populations. Light blue areas indicate the maximum extent of the LGM glaciation (∼21 ka; Ehlers *et al*., 2011). **A**. Distribution of populations with elevated private allelic richness (reflected by dot size) as an indicator for long-term stability of a population and as indication for a refugial area (Widmer & Lexer, 2001). **B**. Number and geographic distribution of genetic clusters (multiple geographically structured clusters vs. single uniform cluster). **C**. Branching of phylogenetic networks (distinct branching reflecting geography vs. shallow or no branching). **D**. Age of divergence between distinct groups or lineages (pre-LGM splits vs. post LGM splits). **E**. Number and location of areas providing suitable climatic niches under LGM climatic conditions (multiple, and separated suitable areas vs. a single suitable area).

Here, we combine information from genomic data and ENMs, all obtained from a range-wide population sampling of three FUS (*Aposeris foetida, Cardamine trifolia* and *Hacquetia epipactis*) with a strong preference for forests dominated by European beech. Their ranges, although overlapping with previously proposed forest refugia, are discontinuous and much smaller than those of beech, suggesting limited dispersal capabilities (Willner et al., 2009). After establishing that the study species constitute a suitable proxy for temperate forests using climate and vegetation data, we asked the following questions. (1) Do the disjunct current distributions of FUS result from long-term survival in multiple glacial refugia followed by limited dispersal to adjacent areas or rather from post-LGM colonisation from a single refugium? (2) Do the identified refugia overlap across FUS in spite of their divergent ranges, and do they match areas that have previously been inferred as temperate forest refugia?

## Materials and Methods

### Study species

Three entomophilic species with strong preference for deciduous forests over carbonate bedrock usually dominated by European beech were studied. *Aposeris foetida* (Asteraceae; distribution area: Fig. 3A) is a hemicryptophyte rosette plant; clonal spread is negligible and there is no adaptation to wind dispersal (Abs, 1994; Chytry et al., 2021). The species is distributed from the colline to the subalpine belt (40–2,080 m a.s.l; Chytrý *et al*., 2016). *Cardamine trifolia* (Brassicaceae; distribution area: Fig. 3A) is a geophyte with pronounced vegetative reproduction via rhizomatous growth (Chytry et al., 2021) distributed from the colline to the high montane belt (55–1,860 m a.s.l; Chytrý *et al*., 2016). *Hacquetia epipactis* (Apiaceae; distribution area: Fig. 3A) is a geophyte with a short rhizome and thus limited vegetative reproduction (Chytry et al., 2021). It thrives from the colline to the montane belt (50–1,600 m a.s.l.; Chytrý *et al*., 2016).

**Fig. 3.**
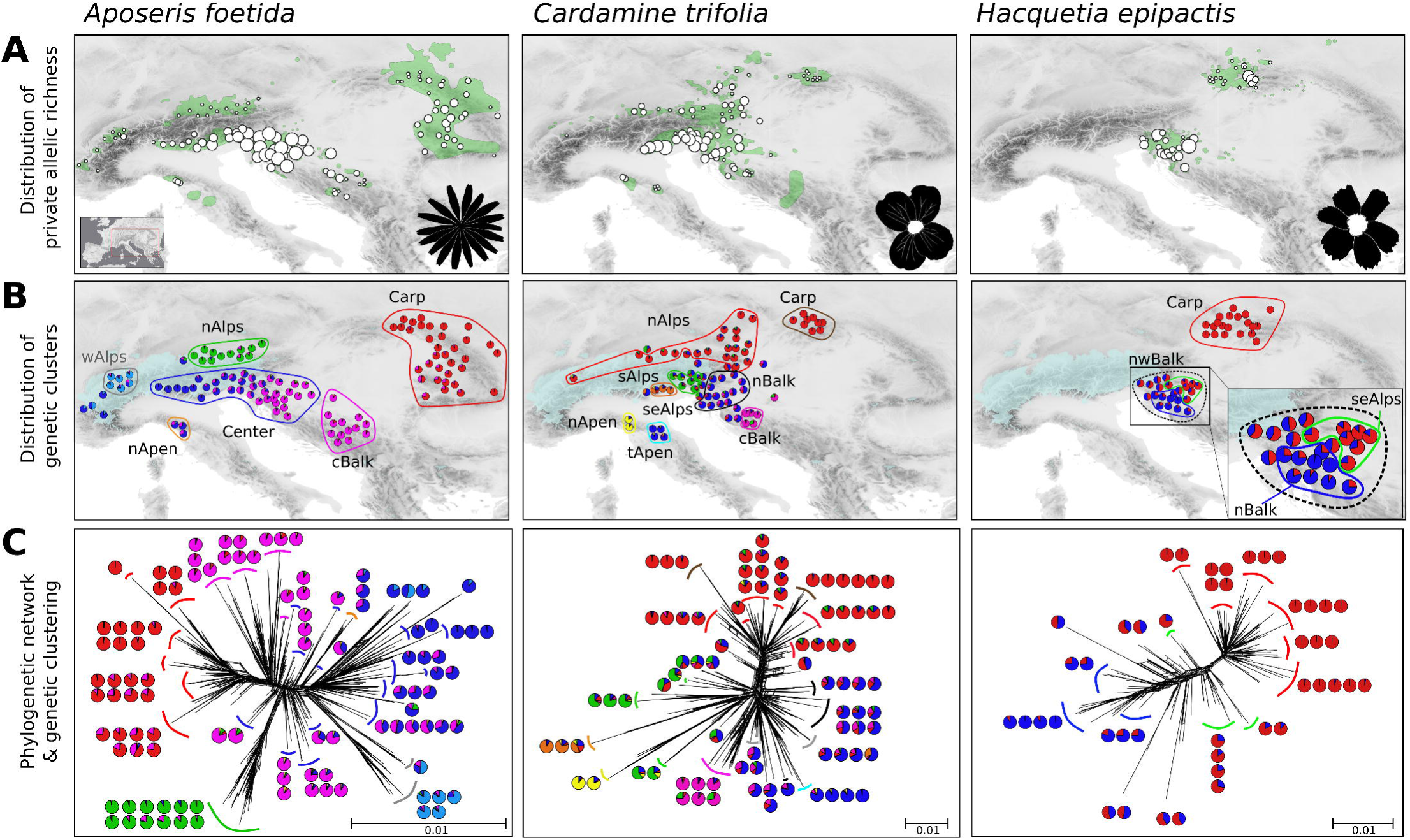
Structure of genetic variation in *Aposeris foetida*, *Cardamine trifolia* and *Hacquetia epipactis* based on RADseq data. **A**. Geographic distribution of private allelic richness. The size of circles reflects the number of private alleles in each population (absolute number of private alleles in Table A1; map including population ID in Supplementary Fig. 1). Green polygons indicate the current distribution of each species (Willner et al., 2023). **B**. Geographic distribution of genetic clusters. Pie charts depict the proportional representation of genetic clusters within each population, as identified by the optimal Bayesian clustering solution (*Aposeris foetida*, K=6; *Cardamine trifolia*, K=6; *Hacquetia epipactis*, K=2). The extent of glaciation at the Last Glacial Maximum (∼21 ka) is indicated in light blue (Ehlers et al., 2011). Colored outlines and labels correspond to geo-genetic groups used in genetic demographic modeling; groups not used are outlined and labelled in light gray. **C**. Phylogenetic networks; colored lines indicate geo-genetic groups as in B. Pie charts show the proportion of genetic clusters in the populations belonging to the respective branches, as identified by genetic clustering.

### Collection of plant material

Samples of *A. foetida*, *C. trifolia* and *H. epipactis* were collected between 2016 and 2019 from 123, 92 and 50 sites, respectively, aiming at covering their entire distribution ranges (Fig. 3A, Supplementary Table 1). Sampling localities (here termed ‘populations’) were numbered from west to east, preceded by species acronyms (Af, Ct and He; Supplementary Fig. 1). From each population, leaves were collected into silica gel from three individuals spaced at least 5 m to avoid sampling of clones; a herbarium voucher was made and deposited in the herbarium of the University of Innsbruck (IB). Sampling was carried out in accordance with the Nagoya Protocol and collection permits were obtained if required (details can be obtained from P. Schönswetter on request).

### DNA extraction

Total genomic DNA was extracted from silica-gel dried material with two different methods. For *A. foetida*, we used a modified CTAB-protocol (Tel-Zur et al., 1999). Prior to extraction with a CTAB buffer the ground tissue was washed three times with a buffer containing sorbitol to remove polysaccharides. DNA extracts were purified using the NucleoSpin gDNA clean-up kit (Macherey-Nagel, Düren, Germany). DNA of *C. trifolia* and *H. epipactis* was extracted using the innuPREP Plant DNA Kit II (Analytik Jena, Jena, Germany) as described by (Kirschner et al., 2022). The DNA concentration of all extracts was estimated using a Qubit 4 fluorometer (ThermoFisher Scientific, Waltham, MA, USA).

### RADseq protocol, identification of RADseq loci and SNP calling

Single-digest RADseq libraries were prepared using the restriction enzyme PstI (New England Biolabs; Ipswich, Masschusetts, USA) and a protocol adapted from (Paun et al., 2016). Briefly, we started with 110 ng DNA per individual and ligated 100 mM P1 adapters to the restricted samples. Shearing by sonication was performed with a M220 Focused-ultrasonicator (Covaris; Woburn, Massachusetts, USA) with settings targeting a size range of 200–800 bp and a mode at 400 bp (peak in power: 50, duty factor 10%, 200 cycles per burst and treatment time 90 s at 20°C). Libraries were sequenced on Illumina HiSeq (Illumina; San Diego, California, USA) at VBCF NGS Unit as 100 bp single-end reads (http://www.vbcf.ac.at/ngs/).

The raw reads were quality filtered and demultiplexed based on individual-specific barcodes using Picard BamIndexDecoder included in the Picard Illumina2bam package (available from https://github.com/wtsinpg/illumina2bam) and the program process_radtags.pl implemented in Stacks 2.3(Catchen et al., 2013; Rochette et al., 2019). The RADseq loci were further assembled, and SNPs were called using the ‘denovo_ map.pl’ pipeline also implemented in Stacks. Denovo_map.pl was run on subsets of the raw data to infer the parameters for an optimal loci yield following(Paris et al., 2017). Consequently, a dataset used for subsequent phylogenetic reconstruction was built using a minimum coverage to identify a stack of 5× (-m 5); a maximum number of differences between two stacks in a locus in each sample of three / six (-M 3 / -M 6) and a maximum number of differences among loci to be considered as orthologous across multiple samples of three / six (-n 3 / -n 6) for *A. foetida* and *C. trifolia /H. epipactis*.

The program *populations* implemented in Stacks 2.3 (Catchen et al., 2013; Rochette et al., 2019) was used to export a set of RADseq loci for the different analyses with the following filtering options: minimum percentage of individuals across populations required to process a locus of 80% / 70% (-R 0.8 / -R 0.7) for *A. foetida* / *C. trifolia* and *H. epipactis*; and maximum observed heterozygosity required to process a nucleotide site at a locus of 65% (-- max-obs-het 0.65). In addition, the --write-single-snp flag was used to select only a single (first) SNP per fragment to avoid linked loci for STRUCTURE (Pritchard et al., 2000) and δaδi (Gutenkunst et al., 2009) analyses. These SNPs were exported in *structure* format (-- structure) for STRUCTURE (Pritchard et al., 2000) analysis and *vcf* format (--vcf) for all other analyses.

The datasets were converted in nexus format for SplitsTree with vcf2phylip(Ortiz, 2019). Finally, for δaδi(Gutenkunst et al., 2009) analyses, SNPs (without filtering for minimum percentage of individuals across populations) were filtered using VCFtools 0.1.16(Danecek et al., 2011) by keeping only sites with depth values over all individuals (DP) between 10 and 50 (--minDP 10; --maxDP 50) and genotypes with a quality above 20 (--minGQ 20).

### Exploratory analyses of SNP data

To estimate the number of private alleles and the nucleotide diversity (π) per population, we used the program *populations* in Stacks 2.3 (Catchen et al., 2013; Rochette et al., 2019). In addition, Neighbour-Nets were produced with SplitsTree 4.17.1 (Huson & Bryant, 2006). To determine the number of genetic clusters and the extent of admixture, we used STRUCTURE 2.3.4 (Pritchard et al., 2000) with ten replicate runs for each K, the number of groups, ranging from 1 to 10, carried out using a burn-in of 20,000 MCMC iterations followed by 200,000 additional iterations. The dataset included 340 individuals and 13,738 loci for *A. foetida*, 232 individuals and 8,438 loci for *C. trifolia* and 127 individuals and 10,808 loci for *H. epipactis*. We summarised the results with CLUMPAK (Kopelman et al., 2015). The optimal K was identified as the K, where the increase in likelihood started to flatten out, the results of replicate runs were similar, and the clusters were non-empty. Additionally, the deltaK criterion was employed, reflecting an abrupt change in likelihood of runs at different K (Evanno et al., 2005).

### Demographic modelling based on genomic data

First, intraspecific genetic groups were defined within each of the three species based on a combination of STRUCTURE clustering, Neighbour-Net networks and geographic distributions (further on termed as ‘geo-genetic groups’). In doing so, only populations for which major cluster assignment in STRUCTURE was > 50%, and assignment to second most frequent cluser was < 35% were included, as the underlying demographic signal in admixed populations may be hidden by gene flow (Chikhi et al., 2010). These criteria were e.g. not fulfilled for the central group of *A. foetida* (see Results). Intraspecific geo-genetic groups were further defined as a set of populations for which at least two of the three following criteria applied: i) the populations belonged to a distinct genetic cluster (from STRUCTURE), ii) the populations formed a divergent branch in the phylogenetic network, and/or iii) the populations were geographically very isolated.

For each species, alternative divergence scenarios were tested between pairs of geo-genetic groups (see Supplementary Table 2 for a list of all pairwise comparisons). This was done using the dadi pipeline 3.1.1 (Portik et al., 2017) that is based on the diffusion approximation method of δaδi that analyses 2D-JSFS(Gutenkunst et al., 2009). Model selection included nine predefined models capturing different divergence scenarios (Charles et al., 2018; Portik et al., 2017; Záveská et al., 2021); graphical representation in Supplementary Fig. 2). The selected models capture changes in population size, migration, and divergence times between groups over one or two epochs. Most importantly, the selected models also allow identification of the mode of divergence, e.g. if it was due to a vicariance event (three models) or a founder event (defined as post-divergence exponential growth of the derived population (Charles et al., 2018). To evaluate directionality of divergence events, the latter models were tested in both directions, meaning that each model was tested twice placing each geo-genetic group as source and founding populations.

The 2D-JSFS were prepared with easySFS (https://github.com/isaacovercast/easySFS) and with the Find-Best-Projections template of (Portik et al., 2017). Down-projecting allows maximising the number of segregating sites by reducing the number of individuals (alleles) in the 2D-JSFS. The numbers of alleles for each down-projection for each group are shown in Supplementary Table 2. Analyses in the dadi pipeline (Portik et al., 2017) were done using the following settings. All models were initially optimised with 3-fold perturbed random starting parameters followed by two rounds of 2-fold and one round of 1-fold perturbation to estimate the log-likelihood of the SFS given the model (folds = 3,2,2,1). For each round, multiple replicates were run (reps = 60,70,70,80) and parameter estimates were used from the best scoring replicate (highest log-likelihood) to seed searches in the following round (maxiters = 10,10,10,15). Across all analyses, optimised parameter sets of each replicate and a multinomial approach were used to estimate the log-likelihood of the 2D-JSFS given the model. Retaining only a single SNP per RADseq locus allows assuming that loci are unlinked, thus the returned log-likelihood values represent the true likelihoods according to (Portik et al., 2017). Model selection was done using the Akaike information criterion (AIC) and Akaike weights (ωi; Burnham & Anderson, 2002). In addition, goodness-of-fit tests were done following (Barratt et al., 2018). This involved running 100 simulations to obtain a distribution of log-likelihood scores. Pearson’s chi-squared test statistic was used to subsequently check whether the empirical values from the original model selection were inside these distributions. Assuming that SNPs were unlinked, these simulations represent parametric bootstraps (Barratt et al., 2018). Simulations were consequently used to obtain confidence intervals for the parameters ‘fraction of the ancestral population’ (‘s’) and ‘divergence time’ (‘T’, ‘T1’ and ‘T2’), which were then transformed into biological estimates following (Gutenkunst et al., 2009). This was done using the mutation rate of *Arabidopsis thaliana* (7e-9 base substitutions per site per generation; Ossowski *et al*., 2010) and generation times (10 years for weakly clonal *A. foetida* and *H. epipactis*; 20 years for strongly clonal *C. trifolia*). To ascertain whether the simulated times were significantly older than the LGM, a one-sided Wilcoxon test was conducted using the R package ‘stats’ (R Core Team, 2023) to compare the distribution of the simulations with a null hypothesis (i.e. lower than LGM) set to 20 ka, which allows obtaining a confidence interval and a pseudomedian value.

### Plastid DNA sequencing

Five plastid DNA regions totalling 6,500, that is *atpB-rbcL* (Piwczyński et al., 2018), *ndhF*– *trnL*, *ndhJ*–*trnT*, *trnH–psbA* and *trnQ*–*trnK* (Shaw et al., 2005, 2007), were inspected for variability in *A. foetida*. For *C. trifolia*, the five before-mentioned plastid regions plus *rpo*B- *trn*C and *trn*G*-trn*S (Shaw et al., 2005, 2007), totalling 8,200 bp, were screened. The *trnH*– *psbA* spacer was most variable and was thus amplified and sequenced for one individual per population from 111 and 83 populations in *A. foetida* and *C. trifolia* (Supplementary Table 1). Amplification was performed in a reaction mix (total volume 20 µl) containing 9 µl REDTaq ReadyMix PCR Reaction Mix with MgCl_2_ (Sigma-Aldrich, St. Louis, MO), 1 µl of the primers (10 µM) trnH and psbA (Shaw et al., 2005) and 1 µl DNA template. Cycling conditions were 5 min at 95°C, 30 cycles of 30 sec at 95°C, 60 sec at 48°C and 60 sec at 72°C, followed by 10 min at 72°C.

For *H. epipactis*, four plastid DNA regions totalling 7,500 bp, that is *ndhJ*–*trnT*, *ndhF*–*trnL*, *trnG*–*trnS* and *trnQ*–*trnK* (Shaw et al., 2005, 2007) were inspected for variability. The *trnQ*– *trnK* region was the only variable and its terminal parts (parts of *trnQ*–*rps16* spacer and *rps16*–*trnK* spacer) were thus amplified and sequenced for one individual per population from 40 and 39 populations, respectively (Supplementary Table 1). Despite several attempts we failed to amplify both the *rps16–trnK* and *trnQ–rps1*6 spacers for individuals of populations He_2, He_19, He_20, He_34 and He_39. Amplification was performed in a reaction mix (total volume 20 µl) containing 7.7 µl REDTaq ReadyMix PCR Reaction Mix with MgCl_2_ (Sigma-Aldrich, St. Louis, MO), 1 µl of both primers (10 µM) trnQ and trnK (Shaw et al., 2007), 1 µl BSA (1 mg/ml) and 1 µl DNA template. Cycling conditions were 5 min at 95°C, 35 cycles of 30 sec at 95°C, 30 sec at 60°C and 4 min at 65°C, followed by 10 min at 65°C. PCR programs were run on Eppendorf 5331 thermocyclers (PE Applied Biosystems, Foster City, CA). PCR products were purified with *E. coli* Exonuclease I and FastAP (Thermosensitive Alkaline Phosphatase; Fermentas, St. Leon-Rot, Germany) following the manufacturer’s instructions. Sequencing was carried out at Eurofins Genomics (Ebersberg, Germany) using the primer psbA for *A. foetida* and *C. trifolia* and the primers trnK and trnQ for *H. epipactis*. Contigs were assembled, edited and sequences aligned using Geneious Pro 5.5.9 (Kearse et al., 2012).

We constructed statistical parsimony networks using PopART (Leigh & Bryant, 2015) applying the TSC network option (Clement et al., 2000). Gaps were treated as fifth character state and indels longer than 1 bp were reduced to single base pair columns allowing those structural mutations to be counted as single base pair mutations only.

### Modelling environmental suitability

Current climatic conditions and those at the LGM were retrieved from the Chelsa Climate database (Karger et al., 2017, 2023). We selected the LGM data based on the Community Climate System Model version 4 (CCSM4; Gent *et al*., 2011) as this model outperforms other global circulation models in Europe (Fordham et al., 2017). We used three bioclimatic variables highly relevant for the occurrence of the modelled species: mean annual temperature (bio1), annual precipitation sum (bio12) and precipitation seasonality (bio15). These climatic variables, which are available with a spatial resolution of 30″, were projected to a grid of 1 × 1 km cell size using the nearest neighbour method and checked to avoid high correlation (|Pearson’s r| < 0.6). As additional environmental variable we used topographical roughness as the standard deviation of the elevations of 100 × 100 m cells (derived from https://www.eea.europa.eu/data-and-maps/data/copernicus-land-monitoring-service-eu-dem) within each 1 × 1 km cell.

We used ENMs relating species occurrences (presences/absences) to the four environmental variables described above to identify suitable areas for each of the three study species under current and LGM climatic conditions. Locations of sampled populations were used as presences, while 100 pseudo-absences were drawn using the ‘sre’ method within the BIOMOD2 package (Thuiller et al., 2024). Thereby, pseudo-absences were selected outside a species’ range envelope constructed by the default setting of BIOMOD2 (i.e., trimming the occurrences along each environmental axis to 95%). The selection of pseudo-absences was repeated ten times. ENMs were parameterized within the BIOMOD2 framework by means of four modelling techniques using their default settings: Generalised Linear Models (GLM), Generalised Additive Models (GAM), Boosted Regression Trees (GBM) and Random Forests (RF). Model runs were replicated three times for each set of pseudo-absences using 80% of the occurrence data for model parameterization and the remaining 20% for model evaluation using the True Skill Statistic score (TSS; Allouche *et al*., 2006). Based on the resulting 120 parameterized models per species (3 replicates × 10 pseudo-absence datasets × 4 modelling techniques), we calculated the environmental suitability as ensemble projections of potential species ranges under current and LGM climatic conditions as weighted (by the TSS value of single models) mean of the projected occurrence probabilities of the single models.

### Niche overlap

Niche overlap between the European beech and each study species was computed following (Broennimann et al., 2012). This comprises the calculation of two-dimensional density distributions (dds) of occurrences for each species and the determination of the niche overlap in this environmental space. First, dds were derived from a PCA based on values of the four environmental variables (standardised to zero mean and unit variance) used for the ENM (including cells all over Europe). Maximum and minimum scores of the first and second axis of the PCA were used to define boundaries of the environmental space, which was divided into 100 bins along each axis resulting in a grid of 10,000 cells each, representing a unique combination of environmental variables. The dd of occurrence points of each lineage was calculated using the function *ecospat.grid.clim.dyn* applying the *adehabit* kernel smoother included in the R package *ecospat* (Di Cola et al., 2017). As a metric for the niche overlap between European beech and each study species, Schöneŕs D (Schoener, 1970) was calculated using the function *ecospat.niche.overlap* from the same package (Di Cola et al., 2017).

### Vegetation analysis

Vegetation data from the European vegetation archive (EVA; Chytrý *et al*., 2016) was used to estimate the amount of variability in the species composition of communities where at least one of the three studied species or European beech were recorded. In total, the data set comprised 19,084 vegetation surveys (i.e., plot records) of 100–200 m^2^, of which 1,417 contained *A. foetida*, 498 *C. trifolia* and 221 *H. epipactis*. A total of 18,051 plots contained European beech. All plot records used were limited to the study area (EVA plot identifiers are provided in Supplementary Data 1). An ordination analysis (detrended correspondence analysis, DCA) was done using the R-package ‘vegan’ (Oksanen et al., 2022).

## Results

### Illumina reads and population structure

Restriction site-associated DNA sequencing (RADseq) datasets resulted in 296,281, 141,559 and 219,077 loci for *A. foetida*, *C. trifolia* and *H. epipactis* after quality filtering, with a mean coverage of 16.0× (±3.2×), 12.3× (±1.0×) and 13.6× (±2.9×), respectively. All resulting raw RADseq data are available in the NCBI Short Archive as BioProject PRJNA824654 (accession numbers SAMN2773504–SAMN27735362, SAMN27409840–SAMN27410080, SAMN28084126–SAMN28084261 for *A. foetida*, *C. trifolia* and *H. epipactis*, respectively; details are given in Supplementary Table 1). The number of private alleles per population ranged from 37 to 991 in *A. foetida*, from 12 to 296 in *C. trifolia* and from 14 to 518 in *H. epipactis* (Supplementary Table 1; Fig. 3A). Nucleotide diversity (π) amounted to 0.014–0.043 in *A. foetida*, 0.054–0.167 in *C. trifolia* and 0.033–0.104 in *H. epipactis* (Supplementary Table 1; Supplementary Fig. 3).

Results from Bayesian clustering analyses for each species are summarised for the respective optimal solution in Fig. 3, and in detail in Supplementary Fig. 4. For *A. foetida*, the genetic clustering analysis resulted in an optimal separation into six clusters (best K = 6) based on 13,738 unlinked single nucleotide polymorphisms (SNPs; Fig. 3B). Six geo-genetic groups were defined following the elaborated rationale, and after excluding few strongly admixed populations (Fig. 3B,C; Supplementary Fig. 4). Despite meeting two criteria, that is geographic coherence and genetic separation of populations, the Center group was not divided into distinct groups due to the extensive admixture cline. This was done to circumvent complications in downstream analyses. For *C. trifolia*, the genetic clustering analysis resulted in an optimal separation into six clusters (best K = 6; Fig. 3B) based on 8,438 unlinked SNPs. After excluding strongly admixed populations, eight geo-genetic groups were defined using the elaborated criteria (Fig. 3B,C, Supplementary Fig. 4). For *H. epipactis*, the genetic clustering resulted in an optimal separation into two genetic groups (best K=2; Fig. 3B) based on 10,808 unlinked SNPs. Following the exclusion of admixed populations, three geo-genetic groups were defined (Fig. 3B,C, Supplementary Fig. 4).

### Demographic history

Quality assessment and parameter estimates of demographic models are provided in Supplementary Table 2. Parameter estimates from goodness-of-fit tests for the best models are given in Supplementary Table 3. In the following, all parameter values correspond to median values inferred by goodness-of-fit tests. In all cases, the increase of the log likelihood values and decrease of variance across replicates, for each model, indicated convergence into the best-ranked model.

For *A. foetida*, founder event models with migration fitted the data best (Fig. 4). When modelling the split between the pairs of geo-genetic groups Af_nAlps/Af_Center, Af_nApen/Af_Center and Af_wAlps/Af_Center, the size of the founding fraction ‘s’ was comparably large (s > 0.2; Supplementary Table 3). Initial analyses did not allow the identification of any ancestral relationships between the two subclusters of Af_Center. To reduce complexity in the modelling, this entity was not divided into further groups. For the colonisation of Af_Carp from Af_cBalk, we found strong support for a founder event involving a small founder population (s = 0.05). Finally, Af_Center was colonised from Af_cBalk via a founder event with considerable population size (s = 0.14).

**Fig. 4.**
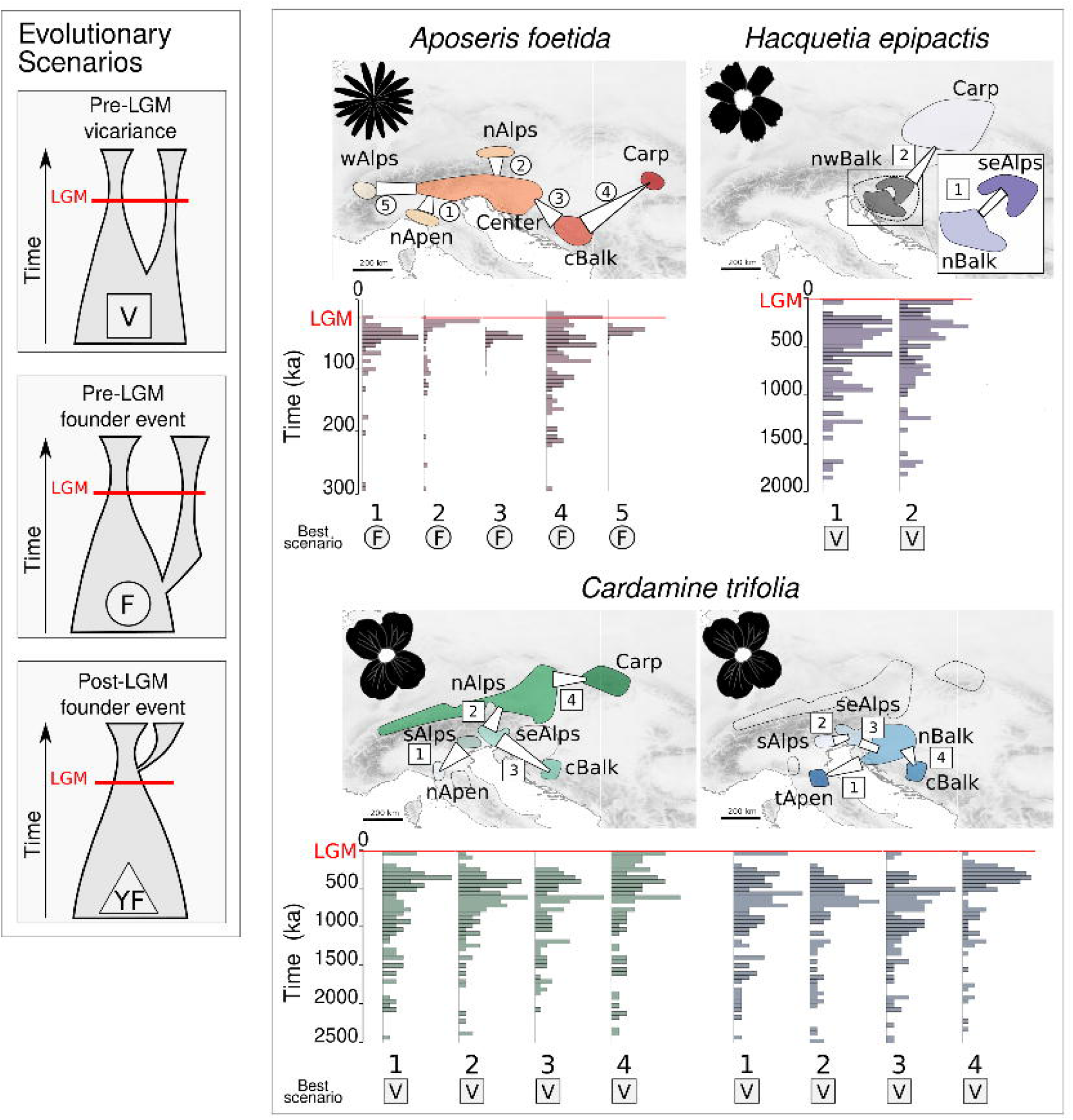
Results of demographic modeling based on genomic data for *Aposeris foetida*, *Hacquetia epipactis* and *Cardamine trifolia*. The panel to the left shows the three evolutionary scenarios that have been evaluated; each scenario is also represented by a symbol (square, circle, triangle) which corresponds with the symbol indicating the best scenario for each species in the right panel. Maps show the geo-genetic groups that were tested against each other; numbers indicate the pairs included in the respective modeling run. The white trapezoids between geo-genetic groups indicate the directionality of the separation whereas the wider side of the trapezoid part faces towards the respective source group. The aspect ratio of sides facing each group proportionally indicates the fraction of the ancestral group involved in establishing the derived group in each modeling run (parameter s; absolute values in Supplementary Table 3). Barplots show the distribution of split age estimates (n = 100) between two groups; numbers correspond to the pairs on the map and the red lines indicate the Last Glacial Maximum (LGM; 21 ka). Results of *C*. *trifolia* are presented in two maps to ensure readability.

All eight pairwise comparisons in *C. trifolia* identified divergence via vicariance followed by secondary contact as the most likely model (Fig. 4). The second-best models received low support in all cases except for the divergence between Ct_nAlps and Ct_Carp (ωi = 0.19).

This model indicated a founder event from Ct_Carp to Ct_nAlps involving a small founder population (s = 0.03), followed by secondary contact. For *H. epipactis* the divergence between He_Carp and He_nwBalk was modelled as vicariance followed by secondary contact. In both best models (ωi = 0.81 and ωi = 0.17, respectively), the split between He_seAlps and He_nBalk was modelled as a pre-LGM vicariance event followed by secondary contact.

For each best-performing model, visualisations of the two-dimensional joint site frequency spectra (2D-JSFS) for empirical data and model data, and corresponding residuals are shown in Supplementary Fig. 5. The results of the goodness-of-fit tests indicated a good fit of all models, and resulted in median divergence times predating the LGM (Supplementary Fig. 5, Supplementary Table 3). One-sided Wilcoxon tests confirmed pre-LGM divergence for all studied pairs of geo-genetic groups (Supplementary Table 3). In some cases, however, the simulated divergence times yielded values post-dating the LGM, e.g. in the pair Af_cBalk/Af_Carp, where 12% of the simulated values indicated post-LGM divergence (Fig. 4, Supplementary Table 2, Supplementary Table 3). In this particular case, the dating result should be interpreted with caution.

### Plastid DNA sequencing

The *trnH*–*psbA* sequences were between 433 (Af_112) and 473 (Af_43) bp long for *A. foetida*, and 313 bp long for *C. trifolia*. For *H. epipactis*, the *trnQ–rps16* and *rps16–trnK* spacer sequences were 843 and 694 bp long, respectively. GenBank accession numbers are given in Supplementary Table 1). The haplotype networks for *A. foetida*, *C. trifolia* and *H. epipactis* identified a central haplotype in each species, that was widespread throughout the distribution range in each species (Supplementary Fig. 6) and present in 75%/83%/81% of the investigated populations, respectively. Divergent haplotypes differed only by one or two mutations, and were distributed across the species’ range as well, with the more common one, aggregated along the Southern and south-eastern Alps in all three species (see Supplementary Fig. 6 for more detail).

### Environmental niche models, niche overlap and vegetation analysis

Regions predicted to be highly suitable by the ENMs were largely consistent across the study species for LGM conditions (Fig. 5) as well as for current day conditions (Supplementary Fig. 7). They comprised large parts of the South and the Southeast of the study area during the LGM, mostly outside of the species current distribution ranges. In contrast, present-day predictions covered well the actual distributions of the species (including all identified refugia) and, in addition, largely matched the current distribution of European beech. All ENMs showed high evaluation scores (TSS = 0.91, 0.93 and 0.93 for *A. foetida, C. trifolia* and *H. epipactis*, respectively).

**Fig. 5.**
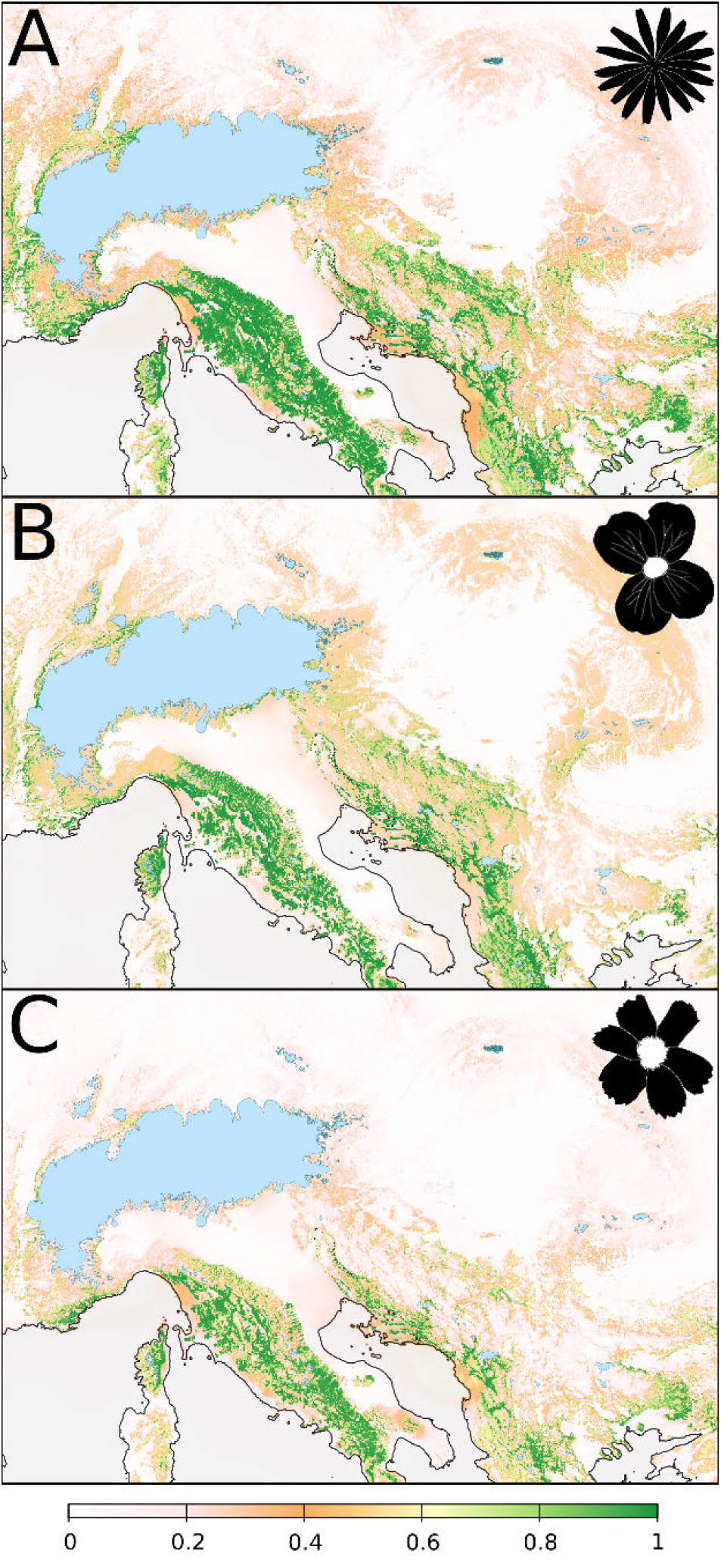
Predicted environmental suitability (indicated by the color shade ramp) for *Aposeris foetida* (A), *Cardamine trifolia* (B) and *Hacquetia epipactis* (C) at the Last Glacial Maximum (∼21 ka; LGM) obtained from environmental niche models (ENM). Coastlines reflect the lower sea level at the LGM (−120 m; Becker *et al*., 2015); LGM glaciation (Ehlers et al., 2011) is indicated in light blue.

The niches of *A. foetida* and *C. trifolia* covered that of European beech to similar proportions, while *H. epipactis* showed a slightly narrower niche (Fig. 6). Each study species rarely occurred outside the niche of European beech. *Aposeris foetida* and *H. epipactis* extended towards higher and lower annual precipitation sums represented by bio12. *Cardamine trifolia* extended towards higher precipitation seasonality and topographic roughness (bio15 and topo). *Hacquetia epipactis* showed two ecological centres of distribution. The empirical niche overlap measured as Schöneŕs D between European beech on the one hand and *A. foetida*, *C. trifolia* and *H. epipactis* on the other hand is D=0.37, D=0.32 and D=0.40, respectively.

**Fig. 6.**
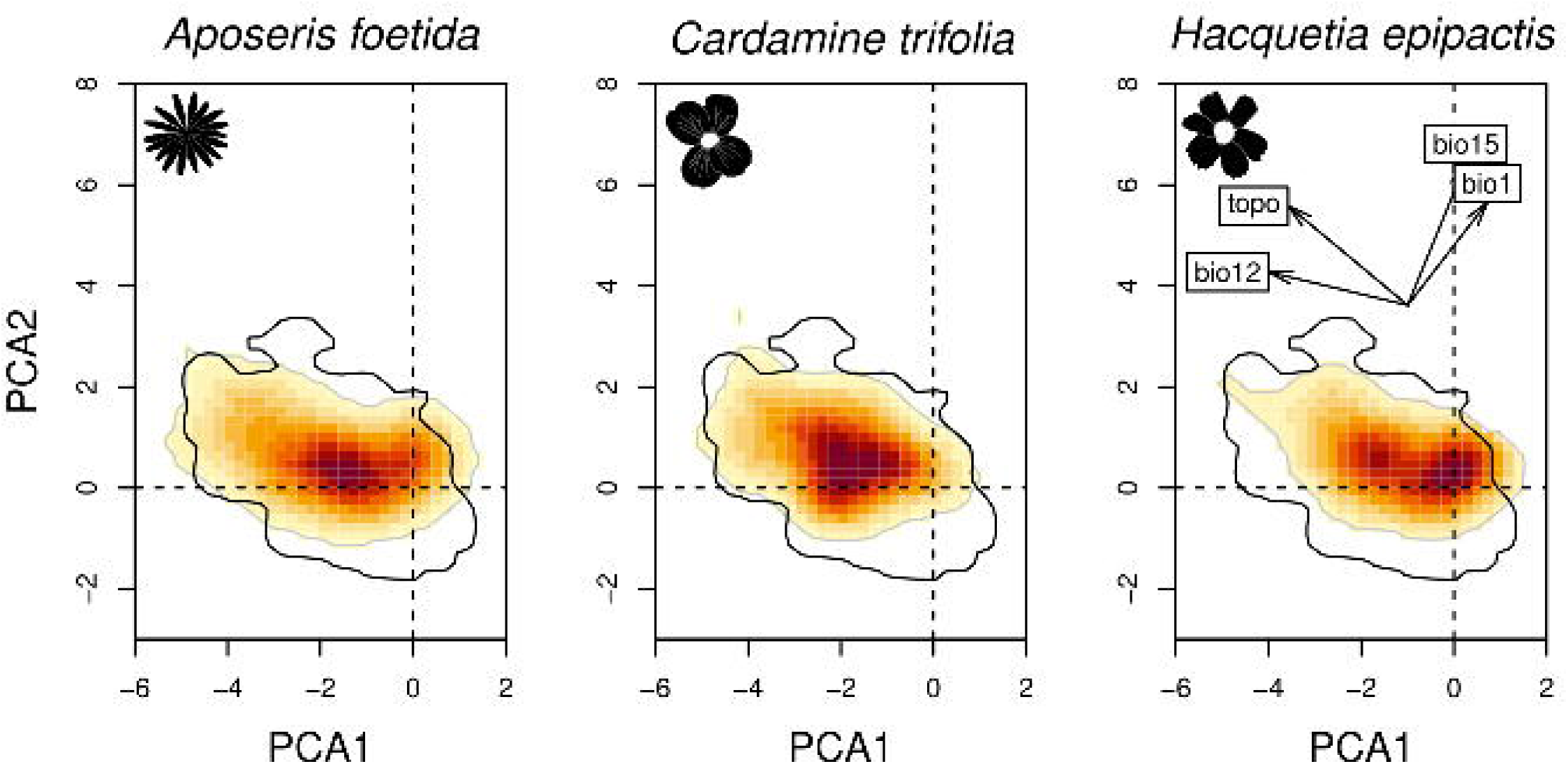
Niche of each forest understorey species (FUS; (shaded area) compared to that of European Beech (*Fagus sylvatica*; contour line) in environmental space (i.e. based on the first and second axis of a PCA using four environmental variables). Shading shows the density of the occurrences of the FUS. A proportion of 37.3% and 30.4% of inertia are explained by the first and second axis of the PCA, respectively. The insert illustrates the contribution of the environmental variables: bio1, annual mean temperature; bio12, annual precipitation; bio15, precipitation seasonality; topo, standard deviation of elevations.

The first and second DCA axis of the vegetation plot records explained 33.2% and 25.9% of variance, respectively. The plot records containing any of the three study species were placed within the variability of records with European beech. *Aposeris foetida* covered most of the ordination space occupied by records with European beech, whereas *C. trifolia* and *H. epipactis* filled a considerably smaller proportion (Supplementary Fig. 8).

## Discussion

The classic hypothesis of Pleistocene cold-stage refugia of European temperate tree species being located in the South of the European peninsulas (Hewitt, 1999; Taberlet et al., 1998) has been challenged by work demonstrating that trees also survived further north in “northern refugia” (Fig. 1). We show that this is also the case in three FUS and outline evolutionary scenarios that involve multiple northern forest refugia in Central Europe. The inferred FUS refugia showed a large spatial overlap between the three species, coinciding well with some of the previously proposed temperate forest refugia (Fig. 1), and also suggesting the presence of forests in areas where this was previously unclear such as the northeasternmost Alps. Among all the identified northern refugia, the northwestern Balkan Peninsula played the most eminent role, further highlighting the status of this region as a diversification centre of European forest biota in general.

We show that the three FUS *Aposeris foetida*, *Cardamine trifolia* and *Hacquetia epipactis* are well-suited proxies for reconstructing the late Quaternary history of Central European temperate forests because their climatic niches are almost entirely restricted to the climatic niche of European beech (Fig. 6) and they occur largely in the same plant communities as European beech (Supplementary Fig. 8). However, some FUS might have been resilient against past changes in the tree composition of temperate forests (Záveská et al., 2021). As a consequence, we suggest that the refugia inferred in this study might indicate the existence of refugia of broadly defined – deciduous, coniferous or mixed – temperate forests.

### The northwestern Balkan Peninsula acted as primary diversification centre

The northwestern Balkan Peninsula and adjacent areas were the primary centre of diversification and acted as the main refugial area for the investigated FUS. As expected for refugia, populations with the highest numbers of private alleles, a proxy for long-term stability (Fig. 3A) and elevated nucleotide diversity (Supplementary Fig. 3) were found in this area. Most of the genetic clusters occurring there exhibit ranges extending beyond the northwestern Balkan Peninsula, which either underlines the region’s role as source for the post-LGM colonisation of other areas or, alternatively, reflects vicariance resulting from past wider continuous distributions (Španiel & Rešetnik, 2022; see following sections). Accordingly, we conclude that this area has acted as a major refugium for temperate forests, which is in line with evidence from recent studies on other FUS (Kirschner et al., 2023; Slovák et al., 2012; Záveská et al., 2021) and on European beech (Fig. 1; Magri *et al*., 2006; Willner *et al*., 2009).

The northwestern Balkan Peninsula was, however, not a single, homogeneous refugium harbouring a panmictic group of refugial populations; rather, it was subdivided and comprised distinct, weakly isolated groups of populations (Fig. 3; Slovák *et al*., 2012). We therefore suggest a scenario where post-LGM expansion from these isolated refugia and subsequent gene flow have caused the weak, but still extant substructure and admixture within the area (Fig. 3B,C). The persistence of genetic structure in the presence of gene flow during post-LGM recolonisation contrasts the assumption that gene flow erases any genetic signal for refugia (Hewitt, 2000; Keppel et al., 2012), but is a prevalent pattern given sufficient resolution of markers (Carnicero et al., 2022; Kolář et al., 2015).

### Survival in multiple northern refugia beyond the northwestern Balkan Peninsula

Beyond the northwestern Balkan Peninsula, northern forest refugia were previously suggested in the Western and Southern Carpathians, the western, northeastern and southern margin of the Alps, the central Balkan Peninsula and/or the northern Apennines (referenced summary in Fig. 1; details in the Supplementary Information). We find refugia linked to almost all disjunct parts of the distribution areas of the studied FUS (Figs. 3, 7). This suggests relative distributional stasis (i.e. no large-scale expansion beyond the refugial range) since the LGM, a phenomenon often observed in species with limited dispersal abilities (Carnicero et al., 2022; Dullinger et al., 2012; Kropf et al., 2002; Schneeweiss & Schönswetter, 2010; Svenning, Normand, & Kageyama, 2008). Indeed, simulations of the postglacial range formation of our study species showed that in the absence of local refugia their disjunct distribution patterns can only be explained by rare long-distance dispersal events (Willner et al., 2023). In addition, demographic models based on genomic data found substantial migration between the tested groups for all three study species (Fig. 4, Supplementary Table 3). However, geo-genetic groups originating from different refugia maintained their genetic integrity over time, even if barriers must have been permeable at different time horizons. This was supported by the best-performing models (Fig. 4, Supplementary Tables 2, 3), and corresponds to evolutionary scenarios outlined for European temperate tree species which similarly exhibited strong genetic structuring despite recurring gene flow (Milesi et al., 2024). In case of *C. trifolia* and *H. epipactis*, almost all models estimated gene flow after an epoch of isolation (Supplementary Fig. 5), which can be interpreted as secondary contact of populations in the course of post-LGM range expansion, which may also support the absence of haplotype structure (Supplementary Fig. 6). We speculated that this indicates ample distributions of the investigated FUS during a warmer and more humid past such as the Holocene climatic optimum (Doláková et al., 2010), when temperate forest thrived also in lowlands such as the Pannonian Plains (the lowlands between the Alps and the Carpathians).

Intraspecific diversification predating the LGM provides strong support for survival in multiple northern refugia. Divergence time estimates derived from the demographic models (Fig. 4, Supplementary Fig. 5, Supplementary Table 3), suggested splits significantly before the LGM (median divergence times: *A. foetida* 40–79 ka, *C. trifolia* 556–897 ka, *H. epipactis* 554–828 ka; Supplementary Table 3; full range of estimates in Fig. 4), providing direct evidence against post-LGM colonisations. In *A. foetida*, however, a small fraction of simulations also resulted in post-LGM splits (12% in nAlps/center; 4% in cBalk/Carp; Fig. 4, Supplementary Table 3, Supplementary Fig. 5), acknowledging that the involved areas (nAlps, cBalk) might not have been refugia. In addition, no explicit demographic test was done between the Southern Limestone Alps and the northwestern Balkan Peninsula in *A. foetida* due to strong admixture of these groups. It thus remains open whether this species also survived in refugia along the southern margin of the Alps.

Divergence events in *C. trifolia* and *H. epipactis* were best represented by demographic models capturing vicariance whereas founder event models represented divergence events in *A. foetida* best. As argued above, this suggests ample, though probably discontinuous distributions of *C. trifolia* and *H. epipactis* during interglacials, which were probably similar to the situation in the Holocene. Such phases were followed by divergence driven by isolation in refugia due to unfavourable climatic conditions during cold stages (Fig. 4, details provided in Supplementary Information). Instead, founder events in *A. foetida* may point to an important role of stochastic dispersal in attaining its present distribution (i.e., long-distance dispersal; Willner *et al*., 2023). However, we emphasise that in the demographic modelling approach used, founder events were defined as an exponential increase of population size of the founded population after divergence (Charles et al., 2018), and consequently, exponential increase of the founding population size may always be resolved as a founder event, irrespective of the initial size of the founding population (s parameter; Supplementary Fig. 2). This contrasts the strict definition of a founder event, which per definition should involve only a small number of individuals (Mayr, 1954; Templeton, 2008). Indeed, in *A. foetida* three out of five models estimated large proportions of the source populations’ effective population size in the founding population (s > 0.2; Supplementary Table 3), hinting at vicariance rather than strict founder events.

### Partially asynchronous divergence and ubiquitous dispersal limitation

Intraspecific splits within *C. trifolia* and *H. epipactis* occurred between ca. 550 ka and 900 ka, significantly predating the LGM (Fig. 4, Supplementary Table 3). As such, these diversification ages are comparable with what has been observed for groups and lineages of other European FUS (Kirschner et al., 2023; Valtueña et al., 2012; Záveská et al., 2021), between Carpathian endemic forest plants and insects, and their sister species (Mráz & Ronikier, 2016), and, recently, also for temperate European tree species (> 600 ka; Marchesini *et al*., 2023; Milesi *et al*., 2024). Notably, in many of the abovementioned instances (including *C*. *trifolia* and *H*. *epipactis*), periods of diversification roughly coincide with the Mid-Pleistocene Transition (0.7–1.25 Ma), when cold stages became significantly longer (ca. 100 ka instead of ca. 40 ka), and more intense (Herbert, 2023), and forests were recurrently replaced by the expanding steppes (Kirschner et al., 2022). In contrast, and as outlined above, divergence within *A. foetida* occurred more recently (Fig. 4, Supplementary Table 3; Supplementary Fig. 6), and was likely shaped by stochastic dispersal events. These younger dispersal events might, however, have erased the signal of older differentiation as has been observed when high numbers of propagules were involved (Todesco et al., 2016). In any event, the lack of synchronicity of divergence events in the study species highlights that codistribution does not necessarily imply congruent evolutionary trajectories (Dawson, 2014). We conclude that the intraspecific diversifications of FUS, although not being synchronous, are much older than the last glacial period, and suggest that instead the Mid-Pleistocene Transition may have significantly shaped the evolution of European temperate forest biota.

Only some of the LGM refugia inferred from genomic data were predicted to be suitable under LGM climatic conditions by ENMs (Fig. 5), while under current climatic conditions, large areas outside of the study species’ actual distribution ranges were found to be suitable (Supplementary Fig. 7). This likely suggests that the fundamental niches of the study species are considerably larger than the currently realised niches (Willner et al., 2023); only the latter are captured by the ENMs. Such discrepancies can be caused by strongly restricted dispersal abilities, preventing the species from reaching areas with climatic conditions that would actually be within their fundamental niche. Indeed, strong dispersal limitation of the studied FUS was suggested by simulations (Willner et al., 2023). It is, for example, very unlikely that the studied FUS inhabited the large areas in Southern Europe predicted as suitable under LGM conditions (Fig. 5), and went completely extinct there after the last cold stage. Most of these regions still harbour deciduous forests similar to those, where the study species occur today (Willner et al., 2009), rendering it likely that the species have actually never reached these areas. Moreover, species might have adapted to the changing climate since the LGM (Lavergne et al., 2010; Luqman et al., 2023; Wessely et al., 2022) which may have contributed to the observed incongruence of highly suitable regions and suggested refugia (Figs. 5, 7).

### Temperate forest refugia in Europe – a synthesis

The refugia suggested for the individual study species (Fig. 7) overlap and mostly match refugia previously suggested for trees based on palynological records and macrofossils (Fig. 1; for an in depth discussion of refugial dynamics for each FUS and each region see the Supplementary Information). Specifically, the northwestern Balkan Peninsula, an unequivocal source of colonisation of Central Europe by European beech (Magri et al., 2006), emerged as a strongly supported refugium based on genomic data from our three study species and additional FUS (Fig. 7; Záveská *et al*., 2021; Kirschner *et al*., 2023). The southwestern fringe of the Western Alps, another well-established refugium (Magri et al., 2006), was supported for *A. foetida*. Refugia at the southern margin of the Alps (Fig. 1; Magri *et al*., 2006; Kaltenrieder *et al*., 2009; Gubler *et al*., 2018) were indicated for *C. trifolia* and *Helleborus niger*, with inconclusive evidence for *A. foetida* and *Cyclamen purpurascens* (Figs. 4, 7; Slovák *et al*., 2012; Záveská *et al*., 2021). The same applies to the northern Apennines, which were identified as a refugium for *C. trifolia* and possibly also for *A. foetida* (Figs. 4, 7).

**Fig. 7.**
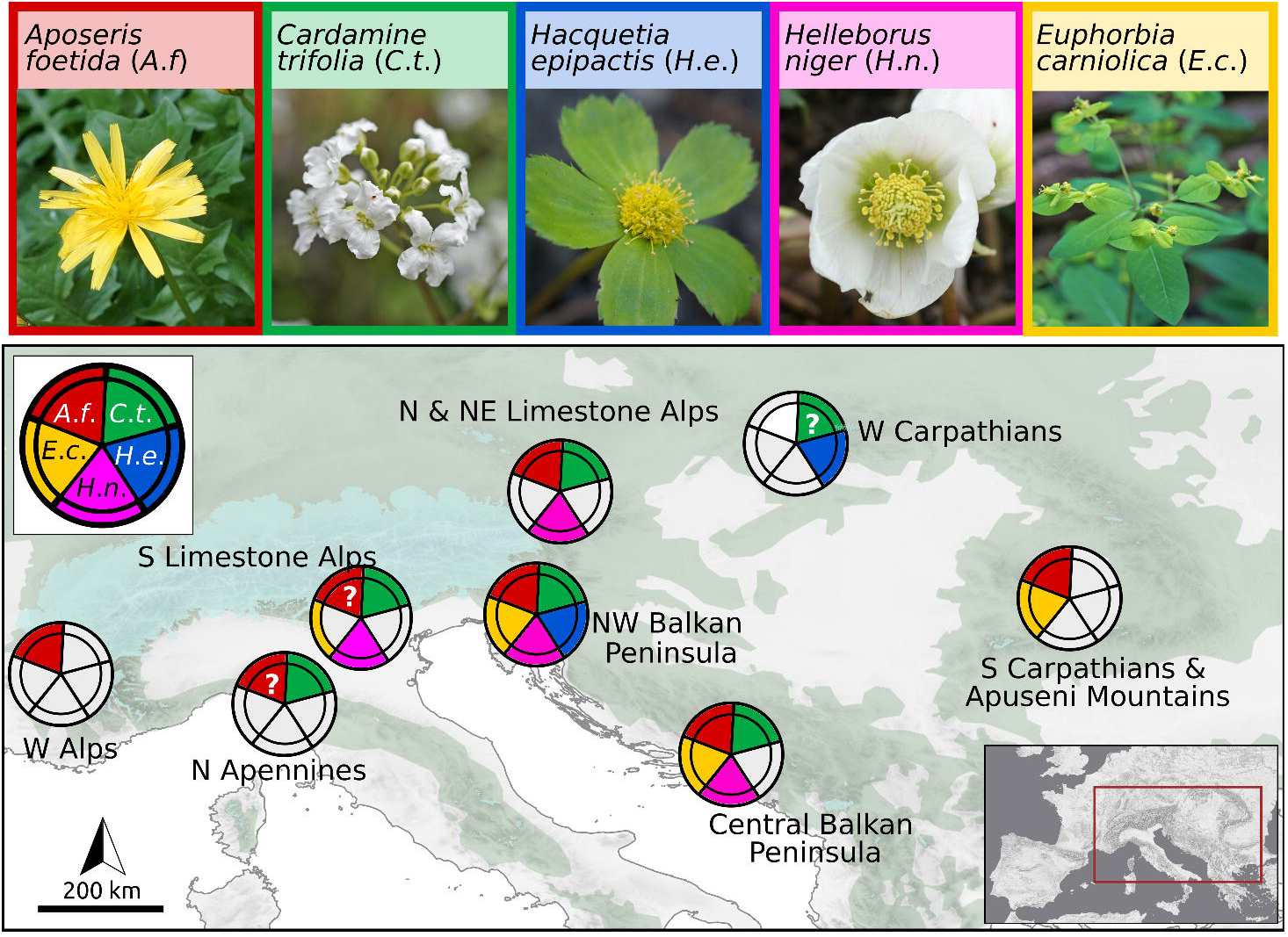
Refugia inferred for understorey species (FUS) of Central European temperate forests based on evidence from genome-wide SNPs (*Aposeris foetida:* ‘*A*.*f*.’*, Cardamine trifolia*: *‘C*.*t*.’, *Hacquetia epipactis*: *‘H*.*e*.’, this study; *E*. *carniolica*: ‘*E*.*c*.’, Kirschner *et al*., 2023; *H*. *niger*: ‘*H*.*n*.’, Záveská *et al*., 2021). Photo credits: Cäcilia Lechner-Pagitz (*A*.*f*., *C*.*t*., *H*.*e*., *H*.*n*.); Božo Frajman (*E*.*c*.). The coloring of the outer tiles and the inner tiles of each circle indicate if a species occurs in the area and if there is genomic evidence for refugial survival of the species, respectively.

Genomic data further support a cold-stage FUS refugium in the Northern Limestone Alps for *A. foetida* and possibly also for *C. trifolia* (Figs. 4, 7). This is remarkable, given that the existence of temperate forests during the LGM has only recently been suggested by genomic data in this region, and so far only in its northeasternmost periphery (Záveská et al., 2021). For the eastern part of the study area, our data support previous findings of glacial refugia (Kirschner et al., 2023; Magri et al., 2006; Marchesini et al., 2023). These refugia have been identified in the Western Carpathians (*H. epipactis* and possibly *C. trifolia*), the Southern Carpathians and the Apuseni Mountains (*A. foetida*, Figs. 1, 7) as well as the central Balkan Peninsula (*A. foetida*, *C. trifolia*). Taken together, our results fully support that the

Northwestern Balkan Peninsula was the preeminent refugium for the investigated FUS; with increasing distance from there, the number of FUS occurring in each potential refugium decreases (Fig. 7). This may suggest that smaller refugia within the area may have limited capacity to preserve a major fraction of temperate forest diversity, similar to the limited carrying capacity of small islands in island biogeography (MacArthur & Wilson, 1967). In any event we emphasise that – disregarding three uncertain cases highlighted in Fig. 7 – all of the disjunct FUS occurrences derive from populations locally present before the LGM, suggesting widespread occurrence of temperate forests through glacial periods.

The existence of widespread northern refugia implies shorter migration distances in recolonizing Europe and explains how temperate forests could have colonised Europe rapidly after the Pleistocene. Based on evidence from FUS, we demonstrate the widespread occurrence of such refugia, and thereby contribute to resolving the long-standing enigma of Reid’s paradox (J. S. Clark et al., 1998). However, in the same line as argued by Záveská *et al*. (2021), our findings hint at the existence of patches of temperate forests in cold-stage conditions rather than providing direct evidence for large-scale deciduous forests in such periods. Some of the outlined refugia may have been dominated by conifers, while still offering suitable conditions for FUS. Indeed, evidence for the existence of different coniferous trees during cold-stages exists for most of the suggested northern refugia or close areas (Fig. 1; Willis & van Andel, 2004; van der Knaap *et al*., 2005; Huber *et al*., 2010; Monegato *et al*., 2015; Gugerli *et al*., 2023). Even under present day conditions *A. foetida* and *C. trifolia* are known to occasionally occur in subalpine dwarf mountain pine (*Pinus mugo*) communities. Finally, we conclude that although FUS refugia do not directly indicate the presence of continuous cold stage forests, they do indicate the presence of trees and the existence of closed canopy cover in all the respective refugial areas outlined in this study.

## Acknowledgments

The paper is dedicated to Christoph Dobeš, who contributed significantly to the research project, from which this paper emerged. He died in a mountain accident in March, 2020. This work was financed by the Austrian Science Fund (FWF, project P29413 ‘Range formation of beech forest understory herbs’ to PS). Marianne Magauer and Daniela Pirkebner performed most of the lab work. Faruk Bogunić, Christoph Dobeš, Julian Haider, Judita Kochjarová, Bohumil Mandák, Ivana Rešetnik, Sabahudin Solaković, Simon Stifter and Ernesto Venturi sampled most of the studied populations. Christoph Dobeš and Iwona Dullinger obtained sampling permits in accordance with national and international laws, and we thank the respective authorities for granting these permits (permits are available upon request). We also thank W. Thuiller and his team (LECA, France) for providing advice in statistics. Analyses were partly conducted using the LEO HPC infrastructure of the University of Innsbruck. Finally, we want to thank the custodians of the European Vegetation Archive (EVA) and its member databases for the permission to use plot data for our analysis.

## Competing interests

The authors declare no competing interests.

## Author Contributions

P.C., P.K., and P.S. designed the study. E.Z. generated the genomic data. C.V., P.C. and P.K. analyzed genomic data. J.W., K.H., and W.W. were responsible for ecological modelling. C.V., K.H., P.C., P.K., P.S., and W.W. co-wrote the manuscript. All authors contributed to the development of the manuscript and improved earlier drafts of the paper.

## Data availability

Short read sequence data generated in this study is available from NCBI short read archive BioProjects PRJNA824654 (accession numbers SAMN2773504–SAMN27735362, SAMN27409840–SAMN27410080, and SAMN28084126–SAMN28084261). Data from chloroplast DNA sequencing was uploaded to NCBI genbank and accession numbers are provided in Supplementary Table 1. Source data for vegetation analyses were obtained from European Vegetation Archive (EVA) and plot identifiers are provided in Supplementary Data 1).

## Notes

### Competing Interest Statement

The authors have declared no competing interest.

