## Supplementary Material1 for "Spatiotemporal diversification of forest understorey species reveals the existence of multiple Pleistocene forest refugia in Central Europe"

**Evolutionary dynamics of forest understorey species in northern refugia**

The **Carpathians** emerged as an area harbouring populations that diverged from other geogenetic groups before the LGM (Fig. 4; Supplementary Table 3), exhibiting little admixture (Fig. 3B,C) and contain private plastid haplotypes (Supplementary Fig. 6A, C). Further, in *A*. *foetida* and *H*. *epipactis* they exhibit an elevated number of private alleles in at least some populations (Fig. 3A), which likely points at their long-term isolation. Within the Carpathians, both the present-day distributions (Fig. 3A; *A. foetida*: Eastern & Southern Carpathians, Apuseni Mountains; *H. epipactis* and *C. trifolia*: Western Carpathians) and the position of populations in the phylogenetic network (Fig. 3C; *A. foetida* and *H. epipactis*: well separated clusters; *C. trifolia*: two groups of individuals partly separated by long splits nested within a cluster of populations from the Northern Limestone Alps) suggest species-specific refugia within or at the periphery of this mountain system. Alternatively, a post-LGM origin of the Carpathians populations of *C. trifolia* through long-distance dispersal from the northern Alps should not be completely excluded, given the very low number of private alleles and the broad distribution of the divergence time estimates (Figs. 3A, 4). Overall, our results fit with previous findings that have suggested that the Apuseni Mountains (Magri et al., 2006) and the Western Carpathians have acted as refugia for FUS (*Aquilegia nigricans* ssp*. subscaposa*, Gafta *et al.*, 2006; *Cyclamen fatrense*, Kučera *et al.*, 2013; *Rosa pendulina*, Daneck *et al.*, 2016; *Syringa josikaea*, Lendvay *et al.* 2016; Mráz & Ronikier, 2016), deciduous tree (Magyari, 2002; Willis & van Andel, 2004; Magri *et al.*, 2006; Birks & Willis, 2008; Juřičková *et al.*, 2014), and insects (Dénes et al., 2016; Drees et al., 2016; Harl et al., 2014; Homburg et al., 2013; Mráz & Ronikier, 2016).

Distinct genetic groups were centred in the **central Balkan Peninsula** in *A. foetida* and *C. trifolia*; whereas in both species the groups’ actual distributions surpassed this area (Fig. 3B). Admixture and the inference of migration with adjacent groups in the northwestern Balkan Peninsula in demographic modeling (Supplementary Fig. 5) suggest extensive and probably recent contact, while the genetic integrity of central Balkan populations has remained largely intact. Together with pre-LGM divergence times (Fig. 4; Supplementary Table 3), we interpret this as evidence in favour of a FUS refugium on the central Balkan Peninsula. This hypothesis fits with several glacial refugia for FUS on the Balkan Peninsula inferred in recent studies (Kirschner et al., 2023). Despite the more or less continuous range of deciduous forests connecting the central and northwestern Balkans under present day climatic conditions, these areas have been separated by steppes and forest steppes during cold stages (Kirschner et al., 2020; Španiel & Rešetnik, 2022), which posed a strong barrier for FUS. The importance of this barrier was underlined in a study on the FUS *Euphorbia* *carniolica* in which the deepest intraspecific split was found between the northwestern and the central Balkan Peninsula, accompanied and most likely also enhanced by ecological divergence (Kirschner et al., 2023). Differentiation in the here studied species was less pronounced, probably due to better dispersal abilities than in myrmecochorous *E. carniolica*.

Northern **Apennine** populations of *A. foetida* and *C. trifolia* (nApen) diverged from geogenetic groups at the southern margin of the Alps before the LGM (Fig. 4; Supplementary Table 3), evidencing the existence of a FUS refugium in the region. The populations were separated by long branches in the NeighbourNet and in case of *C*. *trifolia* they also formed a distinct genetic cluster (nApen; Fig. 3B,C). We interpret this as indication for strong genetic drift due to survival in isolated and small refugia. Interestingly, divergence times of Apennine and Alpine groups of *C*. *trifolia* are similar to divergence times inferred for Apennine and Alpine populations of European beech (Marchesini *et al.*, 2023; Supplementary Table 3). These findings coincide with evidence from distribution patterns (Willner et al., 2009) and palynology (Guido et al., 2020), and explain the presence of forest-margin endemics in this area (Fenu et al., 2022). A further group of Apennine populations of *C. trifolia* (tApen) clusters with those from the northwestern Balkan Peninsula, and originated from a pre-LGM founder event (Fig. 4; Supplementary Table 3). Colonisation may either have involved long-distance dispersal over the Adriatic Sea or stepwise dispersal through the Adriatic Basin, whose northern part dried out during cold stages of the Pleistocene (Becker et al., 2015). Similar scenarios were previously suggested for other plant species or species groups (Falch et al., 2019; Frajman & Schönswetter, 2017; Španiel & Rešetnik, 2022).

In addition to the refugia in the southeastern Alps adjacent to the northwestern Balkan Peninsula, further refugia for FUS were located at the western, northern and southern margins of the Alps. The **northern margin of the Alps** was suggested as refugium for deciduous trees in the past (Juřičková et al., 2014; Tzedakis et al., 2013) but was at the same time cautiously termed a “cryptic refugium” due to the lack of palynological evidence (Tzedakis et al., 2013). We find that populations of *A*. *foetida* and *C*. *trifolia* from the Northern Limestone Alps (in the latter species also from the Bohemian Massif) were clearly divergent from populations in the northwestern Balkan Peninsula (Fig. 3B,C), and diverged before the LGM (Fig. 4; Supplementary Table 3). However, as goodness-of-fit tests for *A. foetida* yielded a small fraction of simulations with post-LGM splits, a recent origin of these populations cannot be fully ruled out (Supplementary Table 3). Overall, our results indicate the existence of a forest refugium in the Northern Limestone Alps, which has been shown for FUS *Helleborus* *niger* (Záveská et al., 2021) and may explain the presence of regional forest understorey endemics such as *Euphorbia saxatilis*, *Callianthemum anemonoides* and *Pulmonaria kerneri* (Essl & Rabitsch, 2009; Kadereit et al., 2019), some of which have undergone a long history of divergent evolution (Frajman & Schönswetter, 2017).

In the **Southern Limestone Alps** two unique genetic clusters were detected in *C. trifolia* (Fig. 3B). The position in the NeighbourNet and the exceptionally large number of private alleles (Fig. 3A,C) both suggest long-term stability and isolation of these populations, and are congruent with the pre-LGM divergence times inferred (Fig. 4; Supplementary Table 3). In *A. foetida*, we found an admixture cline from the Southern Limestone Alps to the northwestern Balkan Peninsula, suggesting secondary contact between previously differentiated populations or a process of differentiation in the absence of complete isolation. We thus suggest the existence of a refugium along the southern margin of the Southern Limestone Alps or perhaps in the Euganean Hills for both species, as has been proposed based on palynological data from broad leaved trees (Gubler et al., 2018; Kaltenrieder et al., 2009), phylogeographic data from other FUS species (*Cyclamen purpurascens*; Slovák *et al.*, 2012); *Rosa pendulina* (Daneck et al., 2016); *Knautia drymeia* (Rešetnik et al., 2016), and by distribution of endemism (Tribsch, 2004).

On the **northwestern margin of the Alps**, pre-LGM divergence times were estimated for *A. foetida* and *C*. *trifolia* (Fig. 4, Supplementary Table 3), providing evidence for isolation in a separate refugium. Additionally, a unique cluster with strong signs of genetic drift was found in *A*. *foetida* (Fig. 3), supporting the refugium hypothesis. This area has not been proposed as a forest refugium before; to our knowledge our study is the first to provide genetic evidence from FUS. Finally, one population of *A. foetida* (Af_11) and one of *C. trifolia* (Ct_3) from the middle part of the Northern Limestone Alps cluster with populations from the southeastern Alps. Several calcicolous species exhibit North-South disjunctions within the Eastern Alps evidencing range expansions over the predominantly siliceous Central Eastern Alps (Merxmüller, 1952, 1953, 1954). Accordingly, glacial survival at the northern border of the Alps, especially in the areas between the major glacier tongues has been invoked to explain these disjunct distribution patterns (Merxmüller, 1952, 1953, 1954), which was later corroborated by phylogeographic studies (Schneeweiss & Schönswetter, 2010). However, recent colonisation events could also explain this genetic pattern, but further analyses with extended sampling are needed to explicitly test this hypothesis.

**Supplementary Figures**

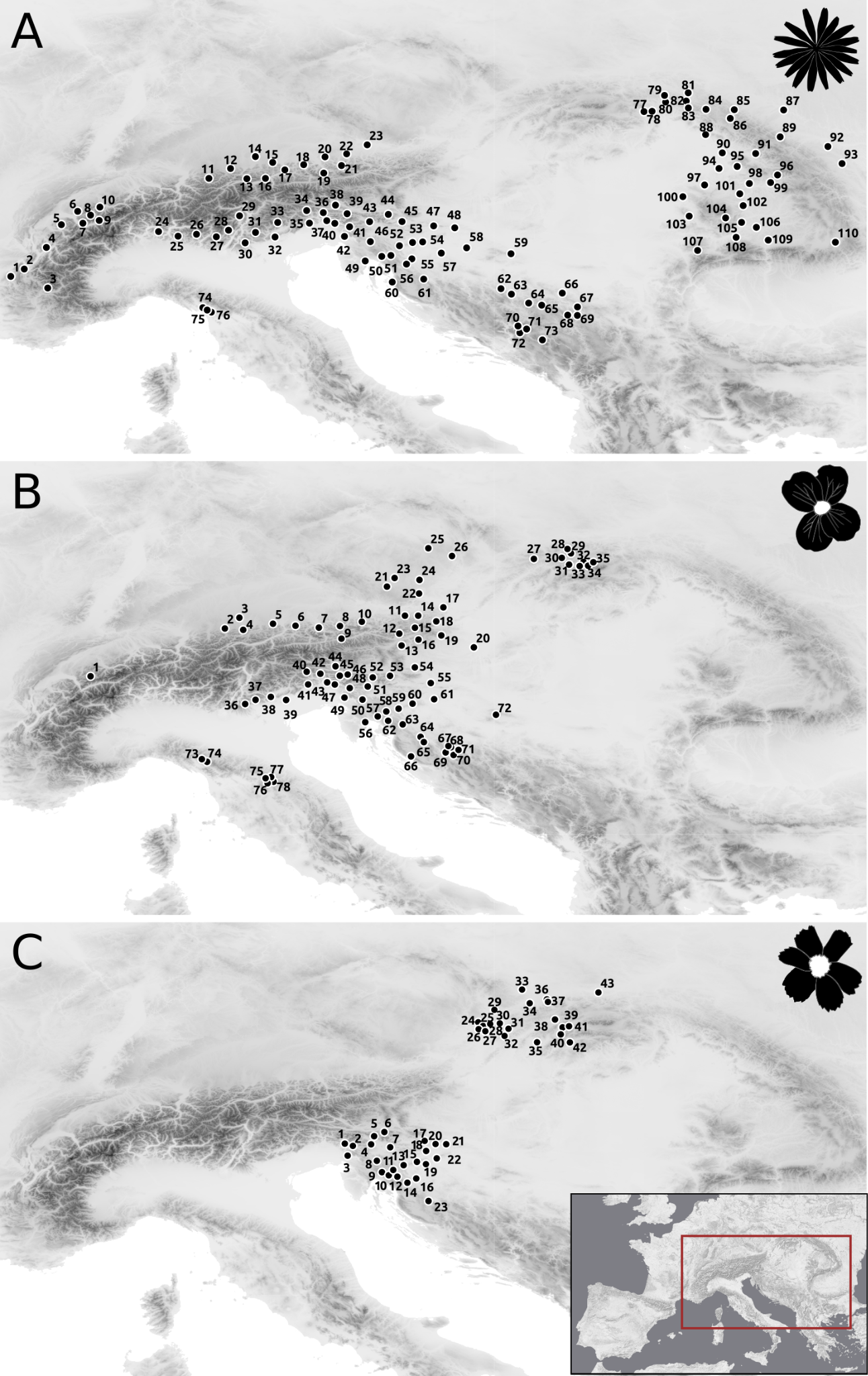

**Supplementary Figure 1.** Sampled populations of *Aposeris foetida* (A), *Cardamine trifolia* (B) and *Hacquetia* *epipactis* (C) as in Fig. 3, but labelled with the population identifiers (details are given in Supplementary Table 1).

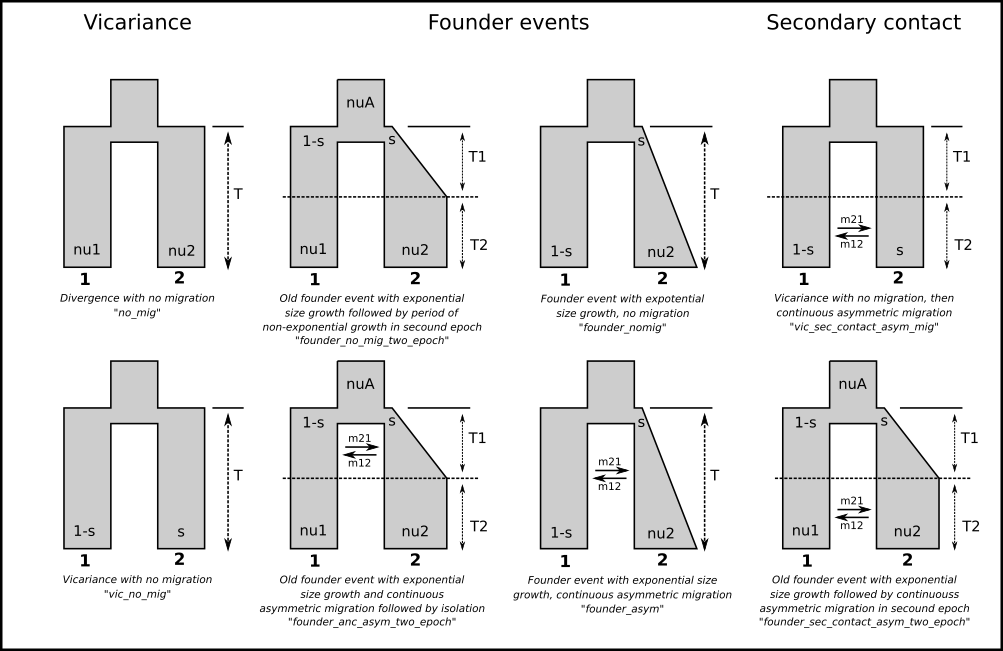

**Supplementary Figure 2.** Graphic representation of the nine models used for genetic demographic modeling. Models were defined as in (Charles et al., 2018; Portik et al., 2017; Záveská et al., 2021).

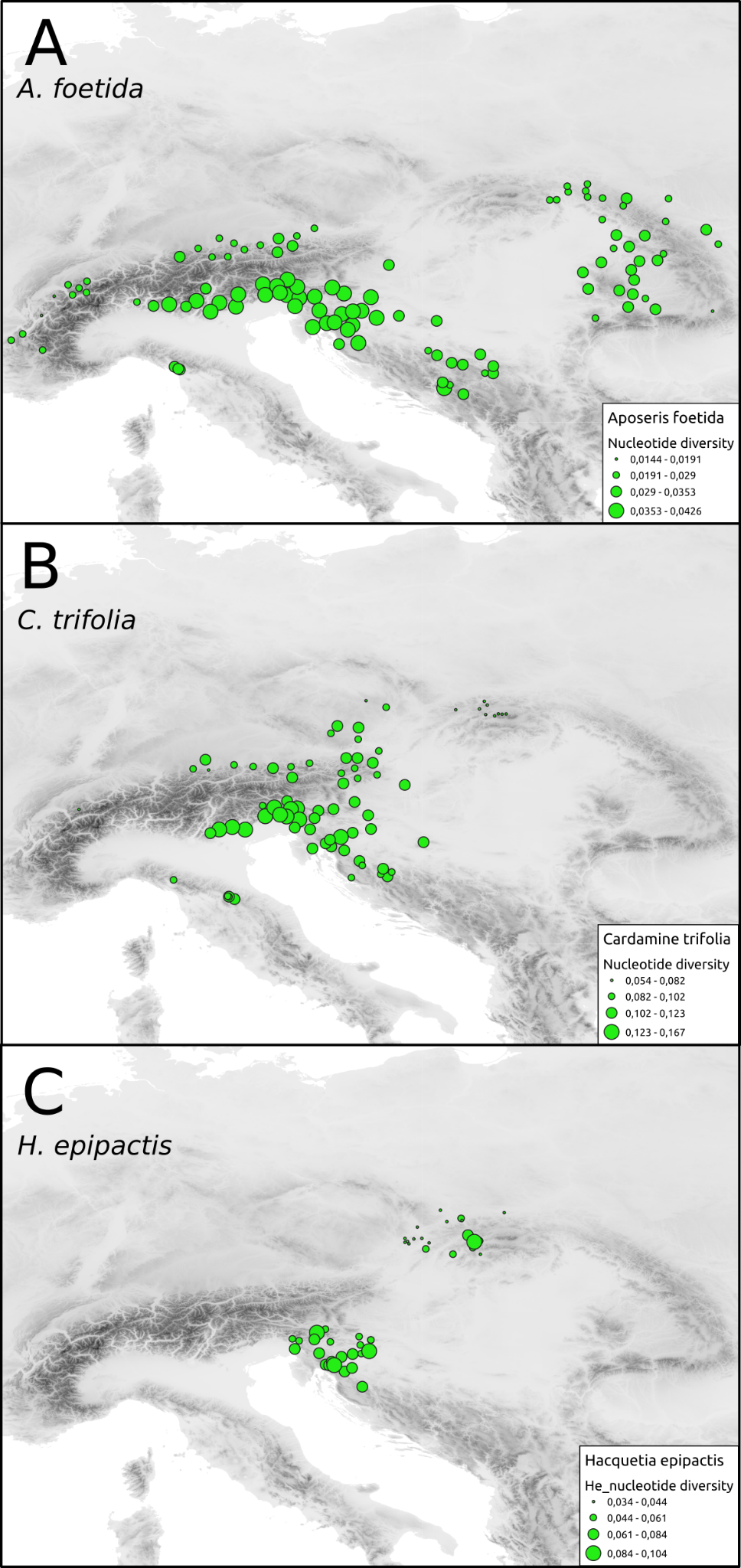

**Supplementary Figure 3.** Extent and distribution of nucleotide diversity for *Aposeris foetida* (A), *Cardamine trifolia* (B) and *Hacquetia epipactis* (C). The size of each circle size reflects the nucleotide diversity in each population, as shown in Table A1.

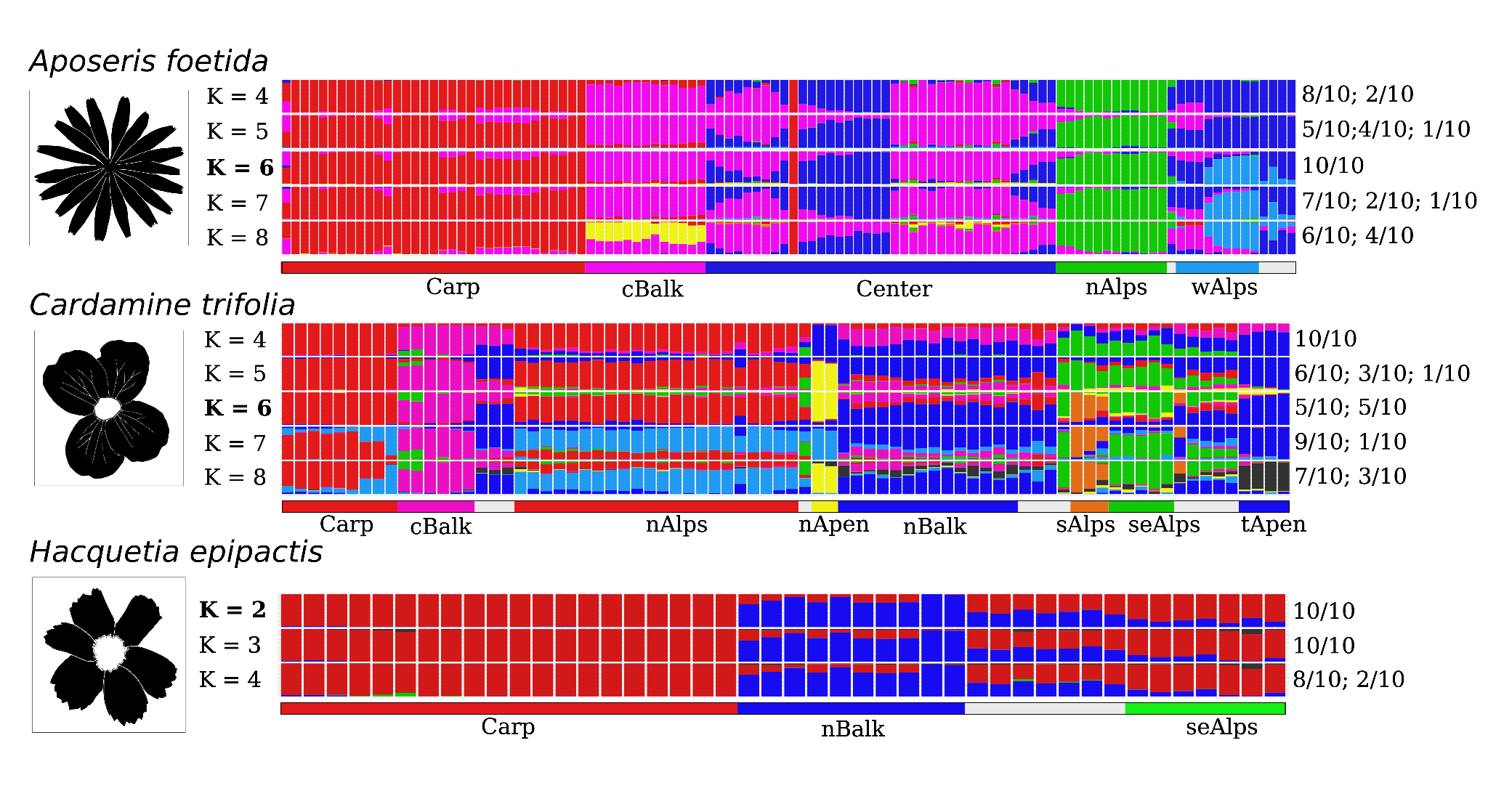

**Supplementary Figure 4.** Results from genetic clustering of *Aposeris* *foetida*, *Cardamine* *trifolia* and *Hacquetia* *epipactis*. Each run of K is shown on the left, with the optimal K (Evanno et al., 2005) being highlighted in bold. Numbers on the right show the proportion of major and minor cluster results (based on 10 replicates for each K). Assignment of each population to the respective geo-genetic group is shown below the barplot for each species. Light grey bars below indicate if populations were removed following the elaborated rationale used for demographic analyses.

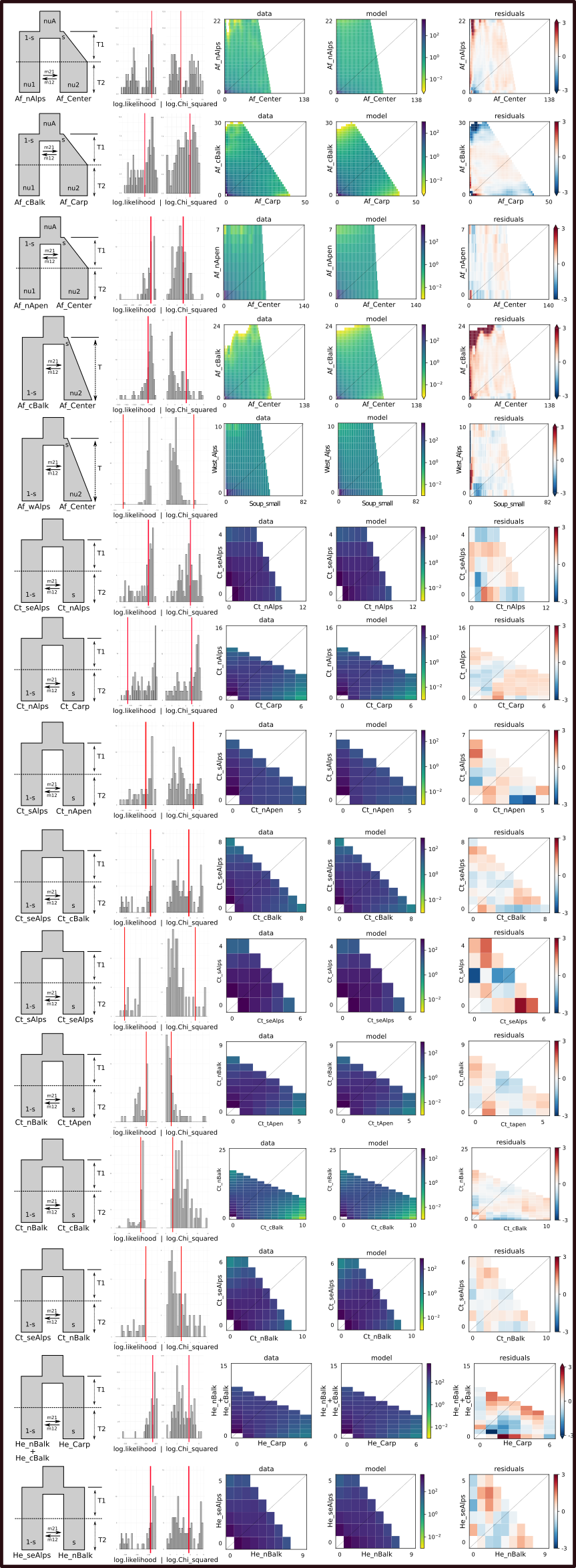

**Supplementary Figure 5.** Best-ranked demographic models selected by δaδi from a choice of nine alternative two-dimensional joint site frequency spectra (2D-JSFS) demographic models for all pairwise comparisons of geo-genetic groups. A visual representation of the best-fit model is depicted, along with comparison of the 2D-JSFS for data, model, and resulting residuals. Next to each model are goodness-of-fit test results, showing the empirical (red lines) value within the distribution of simulation values (grey bars) for log likelihood and Pearson’s log-transformed chi-squared test statistic. Empirical values occurring within distributions indicate good fits. Delimitation of geo-genetic groups as defined in Figure 3.

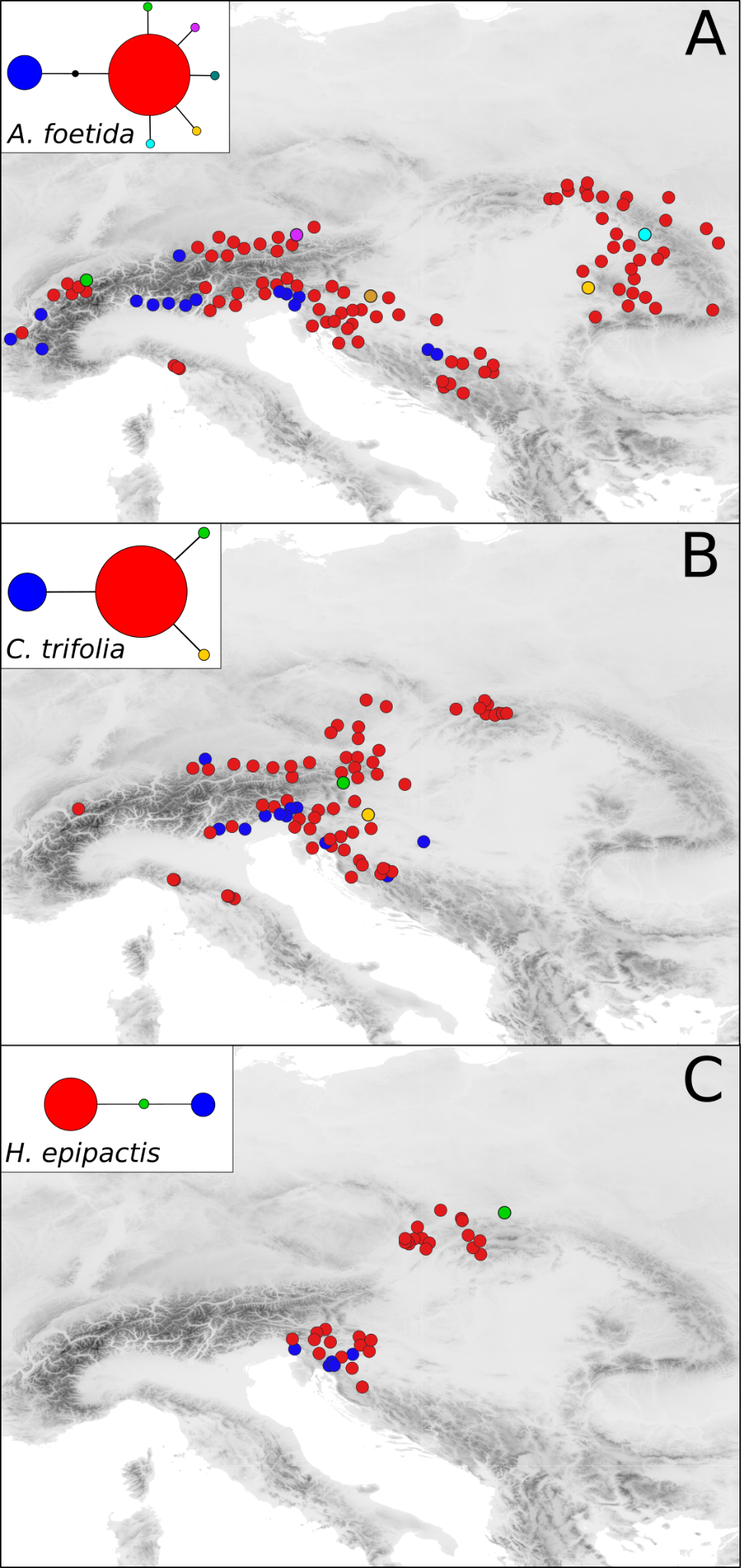

**Supplementary Figure 6.** Plastid DNA variation in the three study species illustrated by statistical parsimony networks (inserts) and maps showing the haplotypes’ distributions. A non-sampled haplotype in *A. foetida* is indicated by a small black dot. *Aposeris foetida* (A) and *Cardamine trifolia* (B) haplotypes obtained from the *trnH*–*psbA* spacer. *Hacquetia epipactis* (C) haplotypes obtained from the *trnQ*–*rps16* and *rps16*–*trnK* spacers. The size of circles in the insert represents the number of populations where the respective haplotype was found.

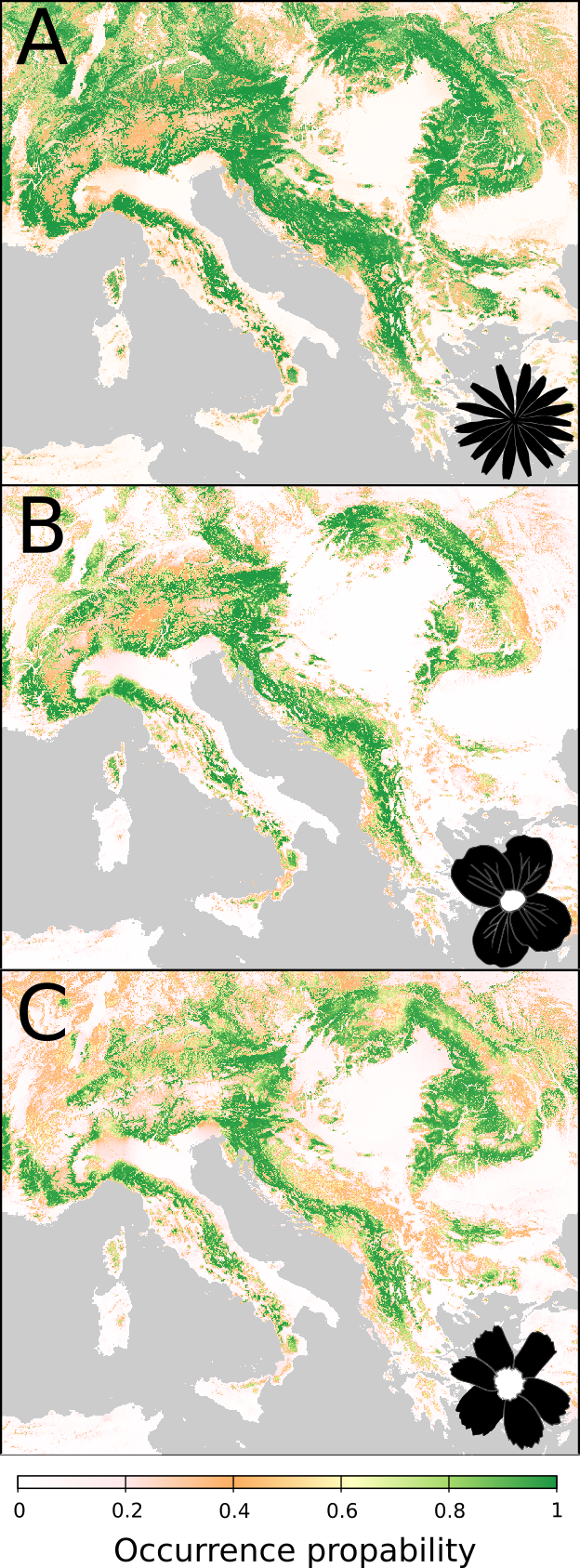

**Supplementary Figure 7**. Predicted present-day occurrence probabilities of *Aposeris foetida* (A), *Cardamine trifolia* (B) and *Hacquetia* *epipactis* (C) obtained from environmental niche models (ENM). The occurrence probabilities derived from ENM are indicated by the color shade ramp below.

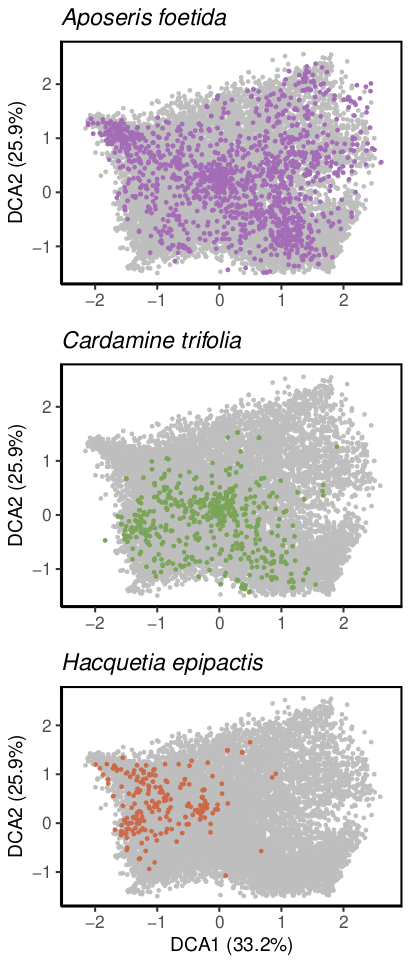

**Supplementary Figure 8.** Detrended correspondence analysis (DCA) of 19,084 vegetation surveys of 100–200 m^2^ containing one of the study species *A. foetida*, *C. trifolia* and *H. epipactis* (coloured dots) and European beech (grey dots). The percentage of explained variation of the plotted DCA axes is indicated in the axis label.
